# Structural and Mechanistic Basis of Substrate Transport by the Multidrug Transporter MRP4

**DOI:** 10.1101/2022.12.12.520055

**Authors:** Magnus Bloch, Isha Raj, Tillmann Pape, Nicholas M.I. Taylor

**Author notes:** Lead contact: Nicholas M. I. Taylor.

## Abstract

Multidrug resistance-associated protein 4 (MRP4) is an ATP-binding cassette (ABC) transporter expressed at multiple tissue barriers where it actively extrudes a wide variety of drug compounds. Overexpression of MRP4 provides resistance to clinically used antineoplastic and antiviral agents, making it a highly attractive therapeutic target for countering multidrug resistance.

Here we report the cryo-EM structures of multiple physiologically relevant states of membrane-embedded human MRP4, including complexes between MRP4 and two widely used chemotherapeutic agents and a complex between MRP4 and its native substrate. The structures display clear similarities and distinct differences in the coordination of these chemically diverse substrates and, in combination with functional and mutational analysis, reveal molecular details of the transport mechanism. Our study provides key insights regarding the basis of the unusually broad substrate specificity of MRP4 and constitutes an important contribution towards a general understanding of the versatile ensemble of multidrug transporters.

## INTRODUCTION

ATP-binding cassette (ABC) transporters are integral membrane proteins that utilize binding and hydrolysis of ATP for moving molecules across cellular membranes. The human genome encodes 48 different ABC transporters that can be divided into seven subfamilies (A-G) based on sequence homology and domain arrangement^1^. Belonging to the category of type IV transporters^2^, subfamily C features a group of efflux transporters characterized by unusually broad substrate specificities^3^ known as multidrug resistance-associated proteins (MRPs).

Unique among the MRPs, MRP4 is found in both apical and basolateral membranes of crucial tissue barriers such as the blood-brain barrier and blood-cerebrospinal fluid barrier^4,5^, where it exports a wide variety of organic anions^6^, including many chemotherapeutic agents^3,7^. In addition, MRP4 has been found to be overexpressed in cancer cells, where its efflux activity contributes to multidrug resistance (MDR): Expression of MRP4 is increased in neuroblastoma^8^, hepatocellular carcinoma^9^, lung cancer^10^, breast cancer^11^, and prostate cancer^12^, and MRP4 overexpression has been found to provide cells with resistance to a wide range of clinically used antineoplastic agents such as thiopurines^13,14^, antifolates^15^, and camptothecins^5,8,16^ as well as enhance the extrusion of antiviral^17–19^ and antibiotic^20,21^ compounds. The extensive tissue distribution and pronounced substrate promiscuity of MRP4 and other MRPs make them important factors to consider when attempting drug delivery^22,23^, and their MDR-inducing overexpression make them attractive therapeutic targets. However, a detailed understanding of the structural and mechanistic basis of their substrate recognition and translocation is currently lacking, and such insight is essential for developing effective modulators specifically targeting the efflux activity of these promiscuous drug transporters.

While recent structural studies have yielded high-resolution molecular structures of multiple members of the ABCB and ABCG subfamilies^24–41^, the MRPs of the ABCC subfamily remain relatively understudied. Recently, structures of bovine MRP1 reconstituted in a detergent-based environment were determined^42–44^, but structures of human MRPs under biologically relevant conditions have so far not been resolved, precluding a deeper understanding of the mechanism of action of this physiologically and therapeutically important group of proteins.

In this study, we used cryo-electron microscopy (cryo-EM) to determine the structure of membrane-embedded human MRP4 and investigate its substrate coordination and translocation. Functional and mutational studies were used to validate the structural models of MRP4, which provide key insights into the structural and mechanistic basis of its substrate transport.

## RESULTS

### The molecular architecture of membrane-embedded MRP4

MRP4 displays a canonical ABC transporter architecture (Fig. 1A) featuring two globular nucleotide binding domains (NBDs) and two helical transmembrane domains (TMDs). The six transmembrane helices of each TMD connect via extracellular loops and intracellular coupling helices, and each NBD harbors a complete set of ABC sequence elements (the ATP-binding cassette). In combination, the two sets make up two composite nucleotide binding sites: a degenerate site, featuring noncanonical sequence elements, and a consensus site, which is strictly canonical and capable of catalyzing ATP hydrolysis.

**Fig. 1:**
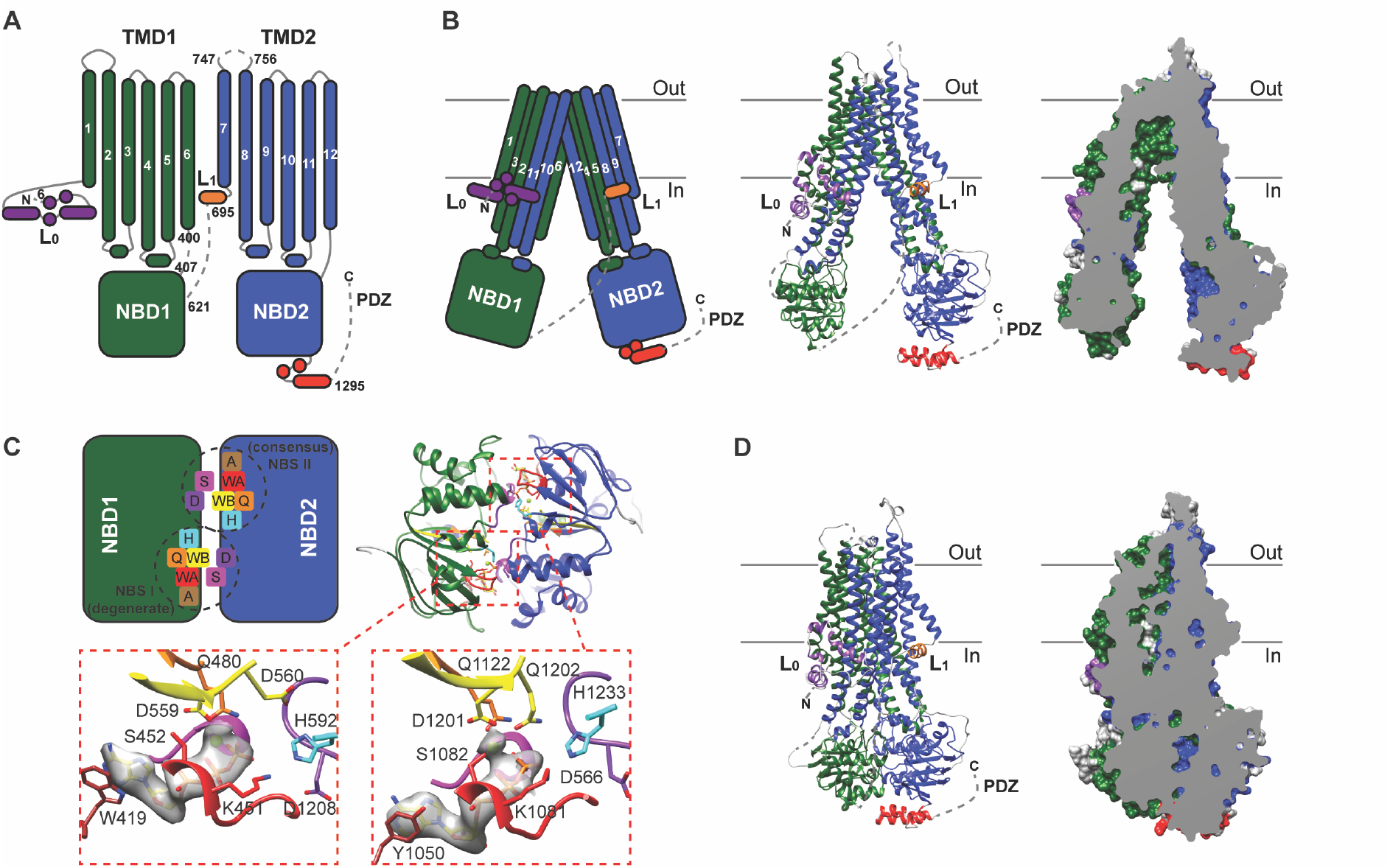
The molecular architecture of human MRP4. A) Cartoon representation of structural elements of hMRP4. Dashed lines represent elements (sequence position indicated with numbers) for which model building was not possible. Transmembrane helices are numbered from N-terminus (N) to C-terminus (C). L_0_ = Lasso motif, L_1_ = C-terminal elbow helix, PDZ = PDZ binding motif. (B) Representations of the hMRP4 IF conformation (viewed from within the membrane plane, indicated with gray lines: Out = extracellular side, In = intracellular side). Left: Cartoon representation of the hMRP4 IF model. Middle: Ribbon representation of the hMRP4 IF model. Right: Slabbed surface representation of the hMRP4 IF model. (C) Representations of the hMRP4 OF conformation (viewed from the intracellular side). Top: Cartoon and ribbon representations of the two composite nucleotide binding sites (NBS I and II) and their ABC motifs (colored boxes): A (brown) = A-loop, WA (red) = Walker A, Q (orange) = Q-loop, WB (yellow) = Walker B, H (cyan) = H-switch, D and S (purple) = D-loop and Signature motif. Bottom: Zoomed in view of isolated ABC motifs. Gray surface represents map density (contoured at σ = 0.70) within 2.0 Å of the molecular models of bound ATP molecules (shown as sticks) coordinated by Mg^2+^-ions (green spheres). (D) Left: Ribbon representation of the hMRP4 OF model. Right: Slabbed surface representation of the hMRP4 OF model.

Like all other type IV exporters^2^, MRP4 features elbow helices immediately preceding the TMDs. The N-terminal elbow helix forms part of an ABCC-specific lasso motif (L_0_) and the C-terminal elbow helix (L_1_) connects indirectly to NBD1 via an extended linker. Among the MRPs, MRP4 features an unusually long C-terminal tail that contains a PDZ binding motif (PDZ) at the C-terminus^45^.

Cryo-EM was used to determine the three-dimensional molecular structure of MRP4. To closely mimic biologically relevant conditions, purified MRP4 was reconstituted in the near-native membrane environment of lipid nanodiscs prior to structural studies.

### The substrate-accepting inward-facing conformation

We were able to reconstitute MRP4 into ‘native’ nanodiscs prepared using brain polar lipid (BPL) extract supplemented with the cholesterol analog cholesteryl hemisuccinate (CHS) (Fig. S1A). From this sample we could reconstruct a map corresponding to a distinct inward-facing (IF) state (Fig. S2).

In this conformation (Fig. 1B), the overall domain arrangement of MRP4 is similar to that observed for other type IV exporters^2^: Two transmembrane helices from each TMD domain-swap into the other TMD forming two helical bundles, with each bundle interacting directly with one NBD via its associated coupling helices. Characteristic of the IF conformation, the extracellular and transmembrane parts of the TMDs form close contacts, while the intracellular parts and the NBDs are completely separated. The TMDs form a transmembrane cavity, which is firmly sealed at the outer bilayer leaflet and opens towards the intracellular side. The elbow helices are situated at the cytoplasmic interface of the inner leaflet in an orientation parallel to the lipid bilayer, and L_0_ extends towards L_1_ on one side of the transporter. The initial stretch of the C-terminal tail adopts a distinct helical fold, while the structure of the extreme C-terminal region (including the PDZ binding motif) and the linker (residues 621-695) remain unresolved.

The cryo-EM map of the native nanodisc complex features multiple distinct densities corresponding to tightly bound molecules of either cholesterol or CHS (Fig. S7). To investigate the effects of these annular lipids, cryo-EM was applied to wildtype human MRP4 reconstituted in ‘minimal’ nanodiscs made using synthetic POPC (Fig. S1B). Although the overall shape of the two reconstructions was the same (Fig. S8A and B), the quality of the map obtained from the sample of minimal nanodisc complexes did not allow for detailed model building.

### The occluded outward-facing conformation

Extensive efforts at resolving additional globally distinct conformations of wildtype human MRP4 reconstituted in native nanodiscs were unsuccessful. In an attempt to promote the population of alternative conformational states, the catalytic glutamate (E1202) of the consensus site Walker B motif was mutated to a glutamine. This mutation effectively eliminates hydrolytic activity without affecting ATP binding^46^ and has previously been successfully employed to obtain structures of NBD-dimerized intermediates of other ABC transporters^24,27,35,43^. However, alternative conformations could still not be distinctly resolved using cryo-EM on samples of E1202Q (EQ) MRP4 in native nanodiscs, and it was not until EQ MRP4 was reconstituted in minimal nanodiscs (Fig. S1C) that a map clearly corresponding to a different conformation was obtained (Fig. S3).

In this conformation, the NBDs are closely associated (Fig. 1C), completing the degenerate and consensus nucleotide binding sites at their interface, which both harbor a Mg^2+^-coordinated ATP molecule. The intracellular parts of the transmembrane bundles are in close contact (Fig. 1D), shutting the intracellular opening of the collapsed transmembrane cavity, which also in this conformation is sealed off towards the extracellular side. Consequently, the resolved structure represents an outward-facing (OF) occluded state.

As opposed to the map corresponding to wildtype human MRP4 in native nanodiscs (Fig. S2), the map of EQ MRP4 in minimal nanodiscs (Fig. S3) does not feature any distinct densities corresponding to annular lipids, despite the maps being of similar resolution.

### MRP4 is differentially modulated by chemically distinct substrates

To investigate the functional significance of the observed annular lipids, the ATPase activities of both types of nanodisc complexes (native and minimal) were quantified (Fig. 2) using two assays (ADP-Glo and PiColorLock). Both ATPase activity assays demonstrate that wildtype MRP4 exhibits a markedly (∼4-fold) higher basal activity when it is reconstituted in native nanodiscs compared to when it is reconstituted in minimal nanodiscs. To control for potential copurification of other ATP decomposing factors, the ATPase activity of equivalent preparations (Fig. S1C) of EQ MRP4 and K1081M (KM) MRP4 (the latter carrying a mutation in the consensus site Walker A motif that interferes with ATP hydrolysis^47^) was also investigated (Fig. S1D), and neither of the control preparations found to exhibit detectable ATPase activity. The same two assays were used to investigate how MRP4 ATPase activity is affected by the presence of substrates. The endogenous substrate, prostaglandin E2 (PGE2), strongly stimulates basal ATPase activity of both types of nanodisc complexes, while the exogenous substrate methotrexate (MTX) only moderately stimulates basal activity. Another exogenous substrate, topotecan (TPT), was found to *decrease* the observed ATPase activity below basal levels for both nanodisc complexes. Therefore, all three substrates appear to interact with functional wildtype protein, although they modulate ATPase activity in different ways.

**Fig. 2:**
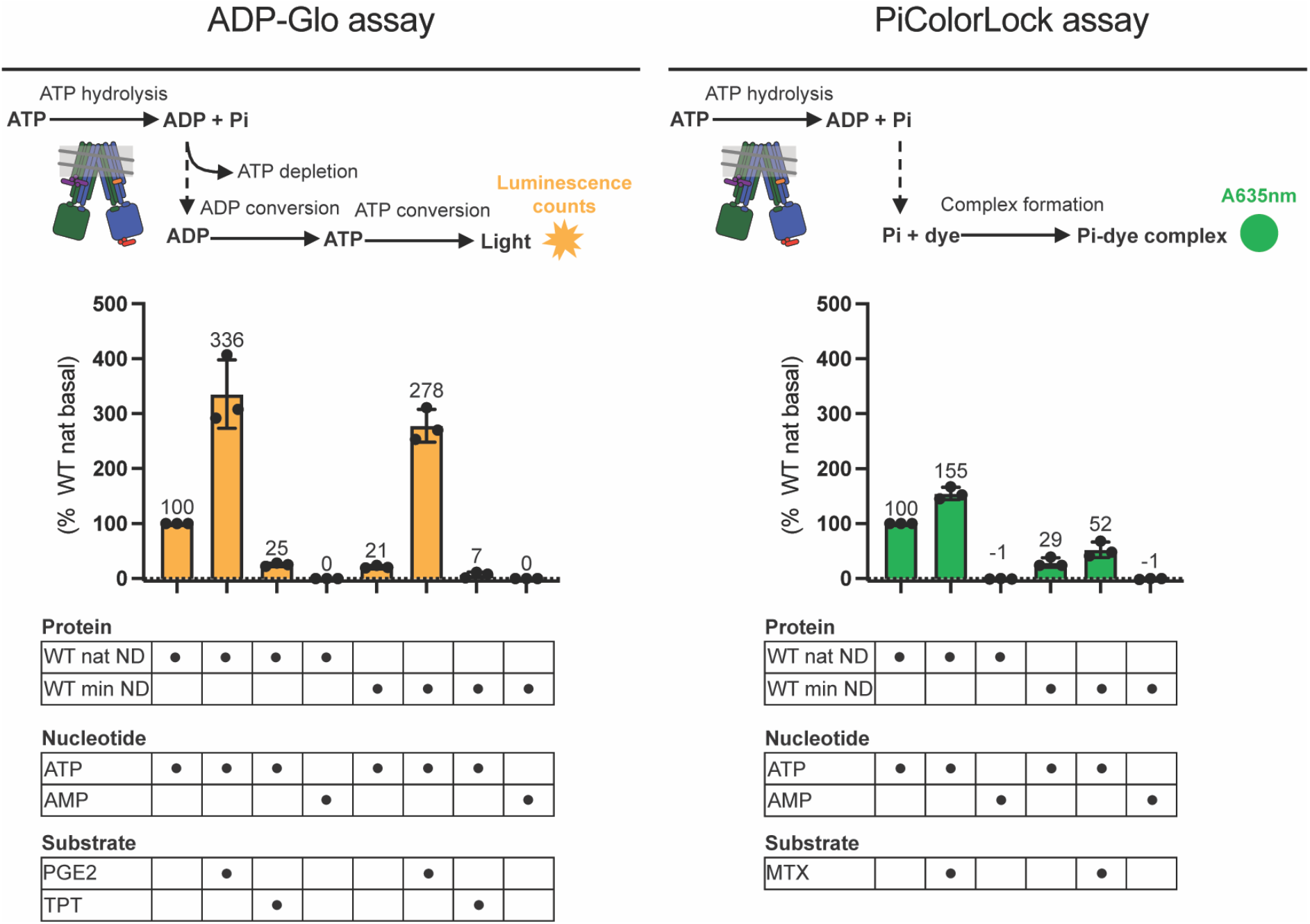
Basal and substrate-modulated ATPase activity of human MRP4. Assays of the ATPase activity of nanodisc-reconstituted hMRP4. Left: Basal and substrate-modulated ATPase activity of wildtype (WT) hMRP4 assayed using the ADP-Glo assay. As shown in the cartoon, ADP-Glo is a bioluminescent assay (with luminescence counts being proportional to the amount of ADP produced from ATP hydrolysis). Right: Basal and substrate-modulated ATPase activity of WT hMRP4 assayed using the PiColorLock assay. As shown in the cartoon, PiColorLock is a colorimetric assay (with the absorbance measured at 635 nm being proportional to the amount of orthophosphate produced from ATP hydrolysis). Activity is normalized to basal activity of WT hMRP4 reconstituted in native nanodiscs (% WT nat basal). Each data point represents an independent experiment (each with three technical replicates) with the average of three independent experiments being reported above each bar (error bars represent the standard deviation). MTX interfered with the ADP-Glo assay (see Method details), which is why the PiColorLock assay was used for samples containing MTX.

### The structural basis of MRP4 substrate promiscuity

To investigate the structural basis of the interaction between MRP4 and these substrates, cryo-EM was applied to samples of WT MRP4 in minimal nanodiscs incubated with either MTX, TPT or PGE2 (see Methods details). The resulting maps (Fig. S4-6) all feature distinct densities inside the transmembrane cavity (Fig. S9A-C) into which molecular models of the respective substrates could be fitted. Consequently, as observed for other ABC transporters, the inward-facing transmembrane cavity appears to constitute a substrate binding pocket (Fig. 3). All three substrates sit in close proximity to the central W995 residue (Fig. S9A-C and 3), which is located at the interface between a predominantly hydrophilic, positively charged half and a predominantly hydrophobic half of the substrate binding pocket, referred to as the P-pocket and the H-pocket respectively^42^. The flat and polycyclic TPT can readily engage in π-π stacking interactions with the indole ring of W995, while the flexible, two-tailed PGE2 potentially engages in H-bonds with multiple residues of the P-pocket via its carboxyl group. The characteristically kinked aromatic moiety of MTX appears to form extensive vdW and π-π stacking interactions with residues of the H-pocket, while the negatively charged carboxyl moiety of the molecule can engage with residues of the P-pocket via H-bonding. Careful examination of the map of IF hMRP4 (no substrate added) (Fig. S2) at low contour levels, revealed a slight density inside the transmembrane cavity (Fig. S9D), which is also observed in a map obtained from a separately prepared sample of WT hMRP4 in native nanodiscs (Fig. S8C and S9E). However, neither were as pronounced as the densities observed in the binding pockets of the structures obtained from the samples with added substrate.

**Fig. 3:**
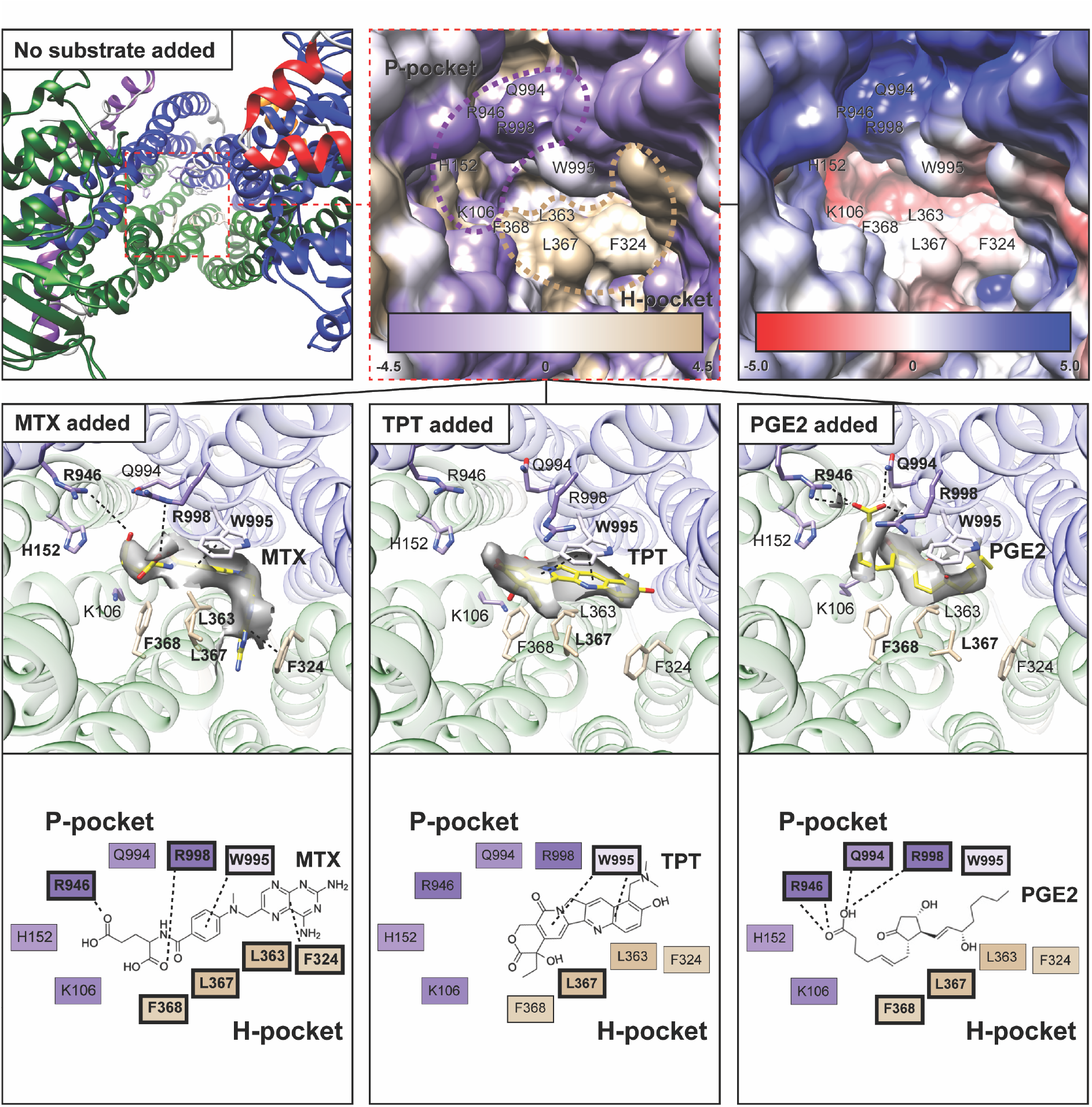
Characterization of the MRP4 substrate binding pocket and its substrate coordination. Top left: Ribbon representation of the molecular model of IF hMRP4 (no substrate added). Top middle: Zoomed in surface representation of the same model with residue surfaces colored according to the Kyte-Doolittle hydrophobicity scale^72^. Scale: purple, negative (−4.5); tan, positive (4.5). Top right: The same surface representation colored according to electrostatic properties. Scale: red, negative (−5.0 kT/e); blue, positive (5.0 kT/e). Middle row: Ribbon representations of molecular models of IF hMRP4 built into the maps obtained from samples to which the indicated substrates have been added (see Method details). Key residues and models of bound substrate are shown as sticks, and potential directed interactions are indicated with punctured lines. Gray surface represents map density within 2.0 Å of the molecular models of MTX, TPT, and PGE2 (contoured at α = 0.40, 0.27, and 0.20 respectively). Residues whose side chains potentially engage directly with bound substrate are highlighted in bold. Bottom row: 2D representations (based on information presented in Fig. S9 and the representations above) of substrate coordination with potentially contacting residues outlined in bold and directed interactions indicated with punctured lines.

### Multiple residues are directly involved in substrate translocation

To validate the structural findings and further characterize the mechanistic basis of MTX transport, mutational analysis of MRP4 was conducted in a cellular setting (Fig. 4A). The MTX efflux capability of multiple variants of MRP4 featuring individual alanine-substitutions of key residues of the substrate binding pocket were investigated by challenging cell lines stably overexpressing these variants with MTX. The susceptibilities of the individual cell lines to the antiproliferative effects of MTX (assumed to be inversely proportional to the MTX efflux capability of the overexpressed MRP4 variant) were evaluated by quantifying cell viability at the different MTX concentrations and fitting dose-response curves to the data (Fig. 4B) to obtain estimates of pIC_50_.

**Fig. 4:**
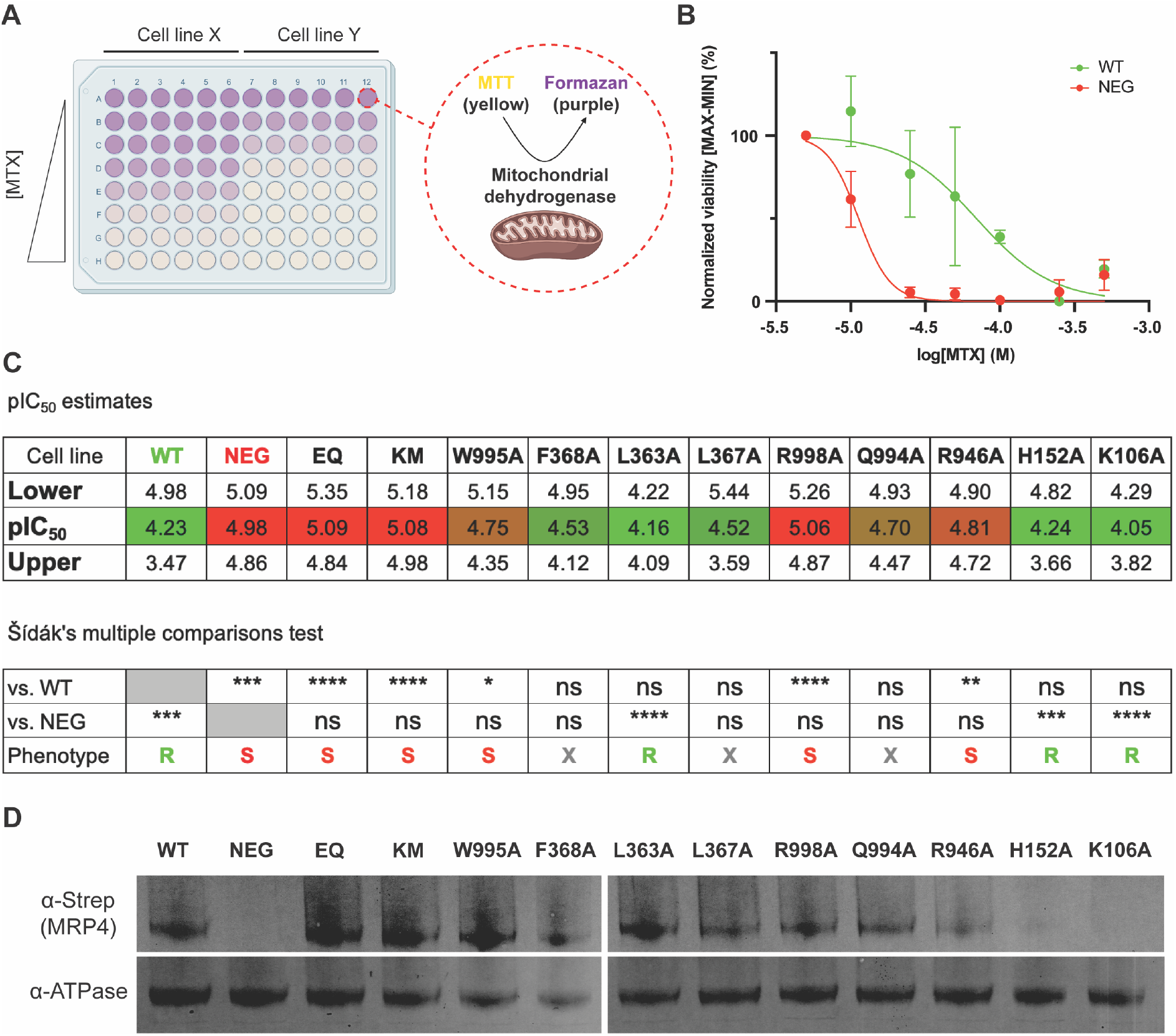
Mutational analysis of the MRP4 substrate translocation mechanism. (A) Cartoon representation of the cellular drug susceptibility assay used for mutational analysis. In principle, the mitochondria of metabolically active cells convert soluble MTT (yellow color) to insoluble Formazan (purple color) with development of purple color thus being proportional to the number of metabolically active cells (and used as a measure of cell viability). (B) Illustrative plot of normalized cell viability vs. MTX concentration (on log_10_ scale) for the cell line overexpressing wildtype hMRP4 (WT) and the parental cell line (NEG). For this plot, normalized cell viability was calculated by normalizing to viability at lowest non-zero [MTX] (5 uM) and then normalizing to the MAX-MIN range. Each data point represents the average of three independent experiments (each with six technical replicates), the error bars represent standard deviations, and the curves represent fits of sigmoidal dose-response models to the data. (C) Table displaying values of mean pIC_50_ (p = -log_10_) for individual cell lines as estimated by the cellular drug susceptibility assay, including Lower and Upper 95% confidence intervals for the estimates (all fits from which these estimates are obtained are shown in Fig. S10). The mean pIC_50_ values are color-coded from red (equivalent to pIC_50_(NEG) = 4.98) to green (equivalent to pIC_50_(WT) = 4.23). The lower table displays the results of a Šídák’s multiple comparisons test and the corresponding phenotypic interpretation (see Quantification and Statistical analysis). (D) Western blots of extracts from equivalent preparations of individual cell lines. Top section: Detection of recombinant hMRP4. Bottom section: Detection of endogenous Sodium Potassium ATPase (loading control).

As expected, the cell line overexpressing wildtype human MRP4 displayed a significantly lower drug susceptibility (see Quantification and Statistical analysis) than the parental cell line (Fig. 4C and S10), which displayed the same phenotype as the cell lines overexpressing the ATPase-deficient E1202Q and K1081M MRP4 variants. Alanine-substitution of the central W995 residue resulted in a sensitive (S) phenotype (Fig. 4C) similar to the one observed for the negative controls. The same phenotype was observed upon mutation of R998 and R946 of the P-pocket, indicating that the identities of these residue positions are critical for normal MRP4 efflux functionality. While phenotyping of the F368A, L367A, and Q994A variants was inconclusive (X), mutations of L363, H152, and K106 yielded resistant (R) phenotypes similar to the one observed for the cell line overexpressing wildtype human MRP4, which indicates that the particular identities of these residues are not of crucial importance for coordination of MTX during transport.

Western blotting was used to assess expression levels of recombinant MRP4 in all investigated cell lines (Fig. 4D). While a complete lack of recombinant MRP4 expression was observed for the parental cell line (NEG), all other cell lines appear to have similar expression levels of their respective MRP4 variant, except for the cell lines overexpressing the H152A and K106A variants. Interestingly, while expression of these variants was not as clearly detected by Western blot analysis, the cell lines nonetheless displayed R phenotypes (Fig. 4C and S10), indicating that they have the same MTX efflux capability as the cell line overexpressing wildtype hMRP4.

## DISCUSSION

### Modulation by annular lipids and substrates

Multiple recent studies have demonstrated the profound effect the reconstitution environment can have on ABC transporter structure and function^48–51^, underscoring the importance of studying these transporters under conditions mimicking their native membrane environment as closely as possible.

In this study, single-particle cryo-EM of human MRP4 in lipid nanodiscs revealed that the TMDs of MRP4 can form specific interactions with lipid molecules that appear to affect conformational dynamics: Native nanodisc complexes of wildtype MRP4, in which ordered lipids associate with the transporter, exhibit markedly higher basal catalytic activity than minimal nanodisc complexes, and while the wildtype transporter preferentially assumes IF states in both lipid environments (regardless of substrate addition), the relatively low resolution of the cryo-EM reconstruction obtained from the sample of minimal nanodisc complex without added substrate may indicate a high inherent flexibility compared to the native complex. Furthermore, only in minimal nanodiscs could the NBD-dimerized (OF) state of EQ MRP4 be resolved, while it remained elusive for equivalent samples of EQ MRP4 in native nanodiscs. Ordered lipids have also been found in recently resolved structures of multiple other ABC transporters^25,37,43^, making it increasingly clear that many if not all ABC transporters form specific interactions with lipids.

However, it is not only the lipid composition of the membrane bilayer that can strongly affect conformational dynamics, but also the presence of substrates and other modulators.

As demonstrated in this study, while presence of substrate modulates the ATPase activity of nanodisc-reconstituted MRP4, the molecular structures of the transporter in complex with substrates (obtained from ‘active turnover’^44,52^ samples, see Methods details) are *globally* indistinguishable from the molecular structures obtained from samples of nanodisc-reconstituted MRP4 to which no substrate has been added (Fig. S11, S2, and S4-6). This contrasts with what is observed from cryo-EM studies of detergent-reconstituted bovine MRP1^42^, where presence of the native substrate, leukotriene C4 (LTC4), appears to induce a distinct relative conformational shift, effectively decreasing the inter-NBD distance (Fig. S11). From recent structural studies^25,42,53^, evidence is accumulating that ligand binding at the substrate binding pockets of ABC exporters affects conformational cycling mainly by altering the relative positioning of the NBDs. Substrate binding may positively modulate ATPase activity by promoting NBD dimerization^25,42^, which would explain the observation that the ATPase activity of most ABC exporters is stimulated by the presence of substrate^54^. However, not all substrates of all exporters stimulate ATP hydrolysis^41^, and drastic, relative NBD-repositioning is not observed in all substrate-bound structures^55^.

### The inward to outward-facing transition

According to the alternating access model^56^, ABC transporters move substrates across cellular membranes by switching between IF and OF states, alternately exposing a central binding pocket to opposite sides of the membrane^57,58^. Canonically, the “resting” state of type IV exporters is the IF conformation^2^, in which the binding pocket is open towards the cytoplasm. Following ATP binding, NBD dimerization causes the exporter to transition into an OF configuration, in which the binding pocket is inaccessible from inside the cell but can be open towards the extracellular side. After ATP hydrolysis, the NBDs dissociate, and the exporter returns to its resting state.

Like other ABC transporters, nanodisc-reconstituted hMRP4 and detergent-reconstituted bMRP1 display significant basal ATPase activity^42^, indicating that both transporters can undergo conformational cycling even in the absence of substrate. Given the widely different reconstitution conditions used for the respective studies of the two different proteins, it is no surprise that they exhibit different conformational landscapes. However, it is nonetheless interesting to observe that nanodisc-reconstituted EQ hMRP4 favors an OF *occluded* conformation, while the equivalent mutant variant of detergent-reconstituted bMRP1 (E1454Q bMRP1) favors an OF *open* conformation^43^ (Fig. S11). Even *wildtype* (detergent-reconstituted) bMRP1 under active turnover conditions^44^ favors the same *open* OF conformation, while wildtype (nanodisc-reconstituted) hMRP4 under similar active turnover conditions is found exclusively in an IF conformation. To what extent this difference in conformational preference reflects the effects of the different reconstitution environments remains to be elucidated.

### The MRP open-occluded isomerization

Neither of the resolved OF structures of hMRP4 and bMRP1 feature bound substrates, despite the same substrates as those of the samples used to obtain the respective substrate-bound structures being present at practically the same concentrations. In the OF occluded conformation of hMRP4, the substrate binding pocket is sealed off and fully collapsed (Fig. S12A), while in the OF open conformation of bMRP1, the pocket is partly accessible from the extracellular side, but rearranged relative to the IF conformations^43^. While both transporters exhibit lateral displacement of their transmembrane helices upon IF-OF transitioning (Fig. S12B), the helix positioning of the occluded conformation most closely resembles an IF configuration. It is therefore likely that the OF occluded conformation of hMRP4 represents a structural state on the MRP transport cycle trajectory closer to transporter resetting (OF-IF transitioning) than the OF open conformation of bMRP1. Interestingly, judging by the relative lateral IF-OF shifts of the hMRP4 and bMRP1 transmembrane helices, open-occluded isomerization appears to involve an asymmetric 8/4 movement of equivalent helices (Fig. S12C) rather than a pseudo-symmetric 6/6 movement of the two transmembrane bundles.

The pronounced relative IF-OF shift of the hMRP4/bMRP1 5/10 helix pair is characterized by strikingly different positionings and orientations of the equivalent binding pocket residues F324/W553: While W553 of OF bMRP1 points *away* from the binding pocket center (and interacts with an ordered CHS molecule), F324 of OF hMRP4 points *towards* the binding pocket center forming a distinct cation-π interaction with R362 (Fig. S13A). A potential equivalent interaction between W553 and R593 of bMRP1 (and between equivalent residues of other MRPs) hints at a common mechanistic feature, and the observation that both W553 and F324 are markedly reoriented outwards in the presence of substrate (Fig. S13B) indicates that this residue position may play an important role in substrate-induced conformational switching of the MRPs in question.

### MRP substrate discrimination

The substrate binding pockets of hMRP4 and bMRP1 share a similar bipartite architecture (Fig. S14A and B), which is reflected in similarities in the substrate profiles of the two homologous transporters: Both exporters preferentially transport organic anions, including metabolic conjugates of GSH, glucuronide and sulfate (e.g. LTC4, E17βG and DHEAS, respectively)^59–62^, and along with other MRPs, both MRP4 and MRP1 also transport MTX^15,63^.

MTX transport by MRP4 has previously been shown to be effectively eliminated by alanine-substitutions of W995 and R998^64,65^, which is confirmed in this study, where also the R946A mutation is shown to significantly reduce MTX efflux by MRP4. Mutation of W1246 and N1245 of hMRP1 (equivalent to W995 and Q994 of hMRP4 and W1245 and N1244 of bMRP1) selectively eliminates E17βG transport without affecting LTC4 transport^66,67^, and charge reversal of R1197 and R1249 of hMRP1 (equivalent to R946 and R998 of hMRP4 and R1196 and R1248 of bMRP1) effectively eliminates not only LTC4 transport, but also transport of E17βG and MTX^68^. Even like-charge substitutions of these residues results in very low MTX transport, and the R998K hMRP4 variant is likewise practically unable to transport MTX^64^, hinting at a general role of these residue positions in MRP substrate discrimination.

While the MTX efflux phenotype of the F368A variant of MRP4 could not be unambiguously determined using the approach of this study, the F368 residue sits at the center of the binding pocket (Fig. 3), and the F368A mutation has previously been shown to effectively eliminate transport of MTX^64^, which indicates that this region is a strong determinant of MRP4 substrate preference. In fact, the neighboring L367 constitutes a unique polypeptide extension among the MRPs (Fig. S14C) and is the only binding pocket residue apart from W995 appearing to be directly involved in coordination of TPT (Fig. 3). These two residues (along with F368) appear to form a hydrophobic sandwich similar to the substrate-coordinating phenyl group sandwich of ABCG2^36–39^, which interestingly enough also binds TPT^41^. While MTX resistance phenotyping of the L367A variant was inconclusive, the exact residue identity of this position may prove crucial for proper transport of TPT and other substrates.

Both MRP4 and MRP1 transport GSH conjugates, which is reflected in the similar spatial arrangement of equivalent P-pocket residues that in bMRP1 coordinate the GSH moiety of LTC4. PGE2 appears to be coordinated by the hMRP4 P-pocket residues similarly to how LTC4 is coordinated by bMRP1 (Fig. S14B), with the two substrates engaging in equivalent interactions with equivalent residues from both transmembrane bundles. However, while LTC4 is a high affinity substrate of both transporters^62,69^, PGE2 is transported with a much higher efficiency by MRP4 compared to other MRPs and does not appear to be a substrate of MRP1^70^. Interestingly, the residues of the hydrophobic sandwich of MRP4 surround the uncharged moiety of PGE2 and may provide a necessary coordination tightness not found in MRP1.

While both MTX and PGE2 appear to interact with residues of both transmembrane bundles of MRP4, their coordination is different (Fig. 3), which may explain why PGE2 strongly stimulates basal ATPase activity, and MTX only moderately stimulates activity. The demonstration in this study that TPT *decreases* ATPase activity of MRP4 to below basal levels, which is also observed for nanodisc-reconstituted ABCG2^41^, may be explained by the observation that TPT does not appear to be as extensively coordinated as the other substrates, and only associates closely with two residues of the substrate binding pocket that nonetheless belong to separate transmembrane bundles.

### The MRP transport cycle

Comparing the obtained structures of MRP4 and MRP1 in light of the accumulated biochemical evidence from functional and mutational analyses of both transporters, we greatly expand our general understanding of the structural and mechanistic basis of the general MRP transport cycle.

While both MRPs display significant basal ATPase activity differentially modulated by the presence of substrates, lipid molecules also appear to affect conformational cycling, reflected in the sensitivity to different reconstitution conditions, and the observation that structures of both transporters feature annular lipids.

The two MRPs exhibit a remarkably similar binding pocket architecture, in which the relatively conserved, positively charged P-pocket section may account for substantial substrate profile overlap, while it appears to be structural variation in other parts of the binding pocket that mainly accounts for substrate profile variation among the MRPs: In MRP4, a tight, hydrophobic sandwich, located at the interface of the two pocket sections, appears to be a strong determinant of MRP4 substrate specificity involved in coordination of both TPT and PGE2, neither of which appear to be substrates of MRP1^70,71^.

The OF occluded conformation of nanodisc-reconstituted hMRP4 presented in this study has to our knowledge never before been resolved for any MRP and likely corresponds to a structural state that must be assumed before transporter resetting can occur^2,52^ (Fig. 5). OF open-occluded isomerization of MRPs practically constitutes opening and closing of a conserved extracellular gate and appears to involve an asymmetric lateral rearrangement of transmembrane helices that for some MRPs may be mechanistically linked to pronounced reorientation of conserved binding pocket residues.

**Fig. 5:**
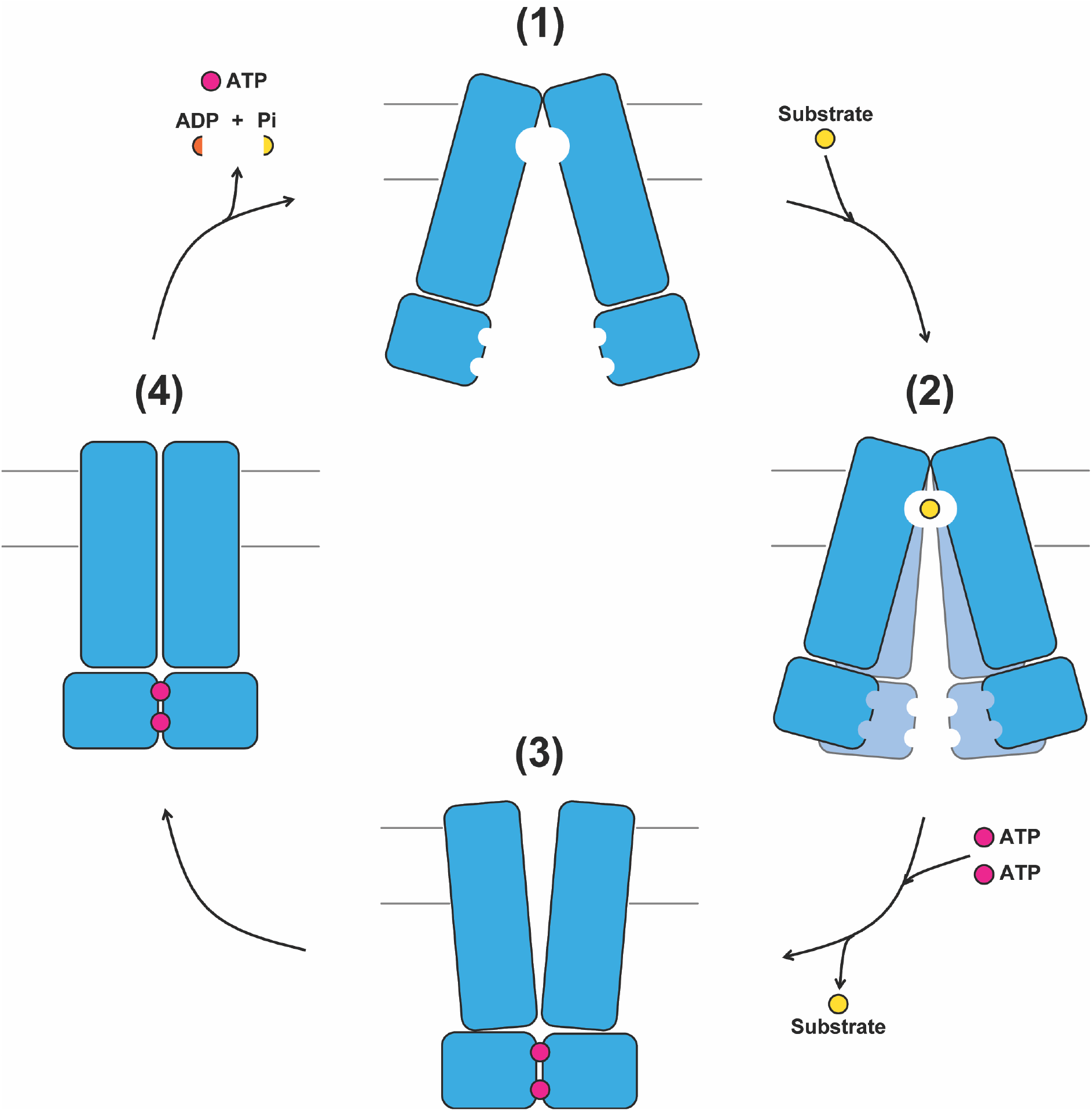
The MRP transport cycle. Cartoon representation of general features of the MRP transport cycle. At least four distinct states have so far been structurally resolved for the MRPs: (1) an IF resting state (for hMRP4 (this study) and bMRP1^42^), (2) an IF substrate-bound state (for hMRP4 (this study) and bMRP1^42^), (3) an OF open state (for bMRP1^43^), and (4) an OF occluded state (for hMRP4 (this study)). While substrate is not necessary for IF-OF transitioning (and its binding does not necessarily effect drastic structural changes), the presence of ATP is apparently required to reach (3), which appears to isomerize with (4). For the asymmetric MRPs (with one nucleotide binding site being degenerate), the (4)-(1) transition (transporter resetting) involves hydrolysis of at least one ATP molecule and release of the produced ADP and orthophosphate (P_i_).

Many details of the conformational trajectory of the MRP transport cycle remain unclear, including the exact mechanism of the allosteric coupling between the binding pocket and the NBDs, and how specifically the interactions between the TMDs and lipid molecules affect conformational cycling. Expectedly, future functional and structural studies will bring us even closer to a general understanding of the physiologically and therapeutically important MRPs.

## ACKNOWLEDGEMENTS

The Novo Nordisk Foundation Center for Protein Research is supported financially by the Novo Nordisk Foundation (grant NNF14CC0001). This work was also supported by an NNF Hallas-Møller Emerging Investigator (NNF17OC0031006). N.M.I.T. is a member of the Integrative Structural Biology Cluster (ISBUC) at the University of Copenhagen. M.B. acknowledges the Neye Foundation for support. We thank the Danish Cryo-EM Facility at the Core Facility for Integrated Microscopy (CFIM) at the University of Copenhagen. Part of the data processing was performed at the Computerome, the Danish National Computer for Life Sciences.

We thank Katharina Duerr (University of Oxford) for kindly supplying us with a vial of the purified pHTBV1.1 vector, Jonathan Elegheert (IINS) for kindly supplying us with the expression plasmid for the HRV3C protease (as well as protocols for expression and purification), and Kenneth A. Matreyek (Case Western Reserve University) for kindly supplying vials of the parental HEK293T LLP-Int-Blast cell line, as well as protocols for maintaining and establishing derivative cell lines.

## AUTHOR CONTRIBUTIONS

M.B. and N.M.I.T. conceptualized the research project and planned the experiments.

M.B. carried out all expressions, purifications and sample preparations for electron microscopy and ATPase activity assays and analyzed the data.

T.P. and M.B. collected and analyzed the electron microscopy data.

I.R. and M.B. built, refined, and validated the structural models.

M.B. conducted the cellular drug susceptibility assays and analyzed the data.

I.R. and M.B. carried out all molecular cloning and generated the derivative stable cell lines.

M.B. wrote the first draft of the paper and prepared all the figures together with N.M.I.T.

All authors contributed to the revision of the manuscript.

## MATERIALS AND METHODS

### RESOURCE AVAILABILITY

#### LEAD CONTACT

Further information and requests for resources and reagents should be directed to and will be fulfilled by the Lead Contact, Nicholas M. I. Taylor.

#### MATERIALS AVAILABILITY

Plasmids generated in this study are available upon request.

#### DATA AND CODE AVAILABILITY

Atomic coordinates for IF MRP4 (no substrate added), OF MRP4 (TPT added), IF MRP4 (MTX added), IF MRP4 (TPT added), and IF MRP4 (PGE2 added) were deposited in the Protein Data Bank under accession codes PDB: 8BJF, 8BWO, 8BWP, 8BWQ, and 8BWR, respectively.

The corresponding electrostatic potential maps were deposited in the Electron Microscopy Data Bank (EMDB) under accession codes EMDB: EMD-16088, EMD-16292, EMD-16293, EMD-16294, and EMD-16295, respectively.

The electrostatic potential maps for IF MRP4 in native nanodiscs (nothing added) and IF MRP4 in minimal nanodiscs (nothing added) were deposited in the EMDB under accession codes EMDB: EMD-16296 and EMD-16297, respectively.

All data reported in this paper will be shared by the lead contact upon request.

This paper does not report original code.

Any additional information required to reanalyze the data reported in this paper is available from the lead contact upon request

### EXPERIMENTAL MODEL AND SUBJECT DETAILS

#### Cell lines

Sf9 cells (used for generating baculovirus) were cultured in Insect-Xpress medium (Lonza, Cat#BELN12-730Q) supplemented with 0.1 % pluronic acid (Gibco, Cat#24040-032) and 50 ug/mL gentamicin (Gibco, Cat#15750-037) at 27°C in suspension.

HEK293-6E cells (used for protein expression) were cultured in expression medium (FreeStyle F17 (Gibco, Cat#A13835-01)) supplemented with 1% FBS (Sigma, Cat#F7524)), 4 mM L-glutamine (Sigma, Cat#G7513), 0.1% pluronic acid, and 50 ug/mL geneticin (Gibco, Cat#10131-027)) at 37°C in suspension.

HEK293T cells (used for drug susceptibility assays) were cultured in plating medium (FreeStyle F17 supplemented with 10% FBS, 4 mM L-glutamine, 1x PenStrep (Sigma, Cat#P4458)), which was supplemented with either 100 ug/mL blasticidin (Sigma, Cat#SBR00022) (for non-transfected cells), 2 ug/mL doxycycline (Sigma, Cat#D3072) (for transfected cells), or 10 ug/mL puromycin (Sigma, Cat#P9620) (for transfected cells) at 37°C in adherent culture.

Frozen vials of the parental cell line (HEK293T LLP-Int-Blast), as well as protocols for maintaining and establishing derivative cell lines, were kindly supplied by Kenneth A. Matreyek (Case Western Reserve University). HEK293T LLP-Int-Blast cells were maintained in T-75 flasks and split every 48-72 hours to a density of 10 M cells/plate.

### METHOD DETAILS

#### Protein expression and purification

The cDNA encoding wildtype human MRP4 was obtained from GenScript (CloneID: OHu17173) and cloned into a donor plasmid based on the pHTBV1.1 vector^73^ to generate a construct featuring full-length MRP4 with a C-terminal HRV 3C-cleavable mVenus-tag linked in tandem to a Twin-Strep-tag and a 6xHis-tag. Mutant variants of MRP4 were generated by divergent PCR with the forward primer carrying the mutated codon. The construct was transformed into DH10Bac cells (Gibco, Cat#10361012) to generate recombinant bacmid (according to the manufacturer’s protocol). Sf9 cells (Thermo Fisher, Cat#11496015) were transfected with purified bacmid to generate baculovirus-infected insect cells (BIICs): 50 ug purified bacmid was incubated for 30 min. with 100 ug Fugene (Promega, Cat#E2311) in 1.5 mL complete Sf9 medium and added to 100 M Sf9 cells resuspended in 30 mL complete Sf9 medium. After 15 min. at 27°C with no shaking, followed by 3 hrs at 27°C with 90 rpm shaking, the culture volume was expanded to 100 mL and incubated at 27°C with 120 rpm shaking for 96 hours before the BIICs were harvested, aliquoted and frozen.

To generate first passage (P0) recombinant baculovirus, 21 M Sf9 cells were allowed to adhere in a T-75 flask for 30 minutes, the medium removed and replaced with 3 mL complete Sf9 medium containing 0.5 M resuspended BIICs. After 3 hrs at 27°C with no shaking, the culture volume was expanded to 11 mL, and the flask left for 96 hours at 27°C with no shaking.

Subsequent passages of recombinant baculovirus were generated in suspension cultures: The P0 culture was harvested, centrifuged (1,500 rcf, 20 min.) and 10 mL supernatant added to a 40 mL 2 M c/mL Sf9 culture, which was left for 72 hours at 27°C with 120 rpm shaking to generate P1 baculovirus. The P2 culture was generated using the same supernatant:culture ratio and harvested by centrifugation. The P2 baculovirus was precipitated by gently mixing P2 culture supernatant with a polyethylene glycol (PEG) solution (20% w/v PEG 10000, 1.2% NaCl) in a 4:1 v/v ratio and leaving it at 8°C with no shaking for > 4 hrs in the dark. Precipitated baculovirus was harvested by centrifugation (3,000 rcf, 45 min.) and resuspended in PBS to 10% of the original supernatant volume to generate a 10x P2 baculovirus suspension.

For expression, the 10x P2 suspension was added at 1% v/v to 2 M c/mL HEK293-6E suspension cultures and the cultures left at 37°C with 150 rpm shaking for 72 hrs. The cells were harvested by centrifugation (500 rcf, 10 min.), the pellets washed with PBS and centrifuged (1,500 rcf, 10 min.) and flash frozen to be stored at -80°C.

For purification, the membrane fraction was isolated: Cells were resuspended in lysis buffer (100 mM HEPES pH 7.5, 200 mM NaCl, 10% glycerol) supplemented with EDTA-free Protease inhibitor cocktail (Roche, Cat#05056489001), 1 mM phenylmethylsulfonylfluoride (Sigma, Cat#93482), and 0.3 mg/mL DNAse I (Sigma, Cat#DN25) and lysed on ice: ∼35 mL aliquots were serially sonicated using nanoprobe. The lysate was cleared by centrifugation (8,000 rcf, 20 min., 4°C), and the membrane fraction isolated by ultracentrifugation (180,000 rcf, 45 min., 4°C) and resuspended at 0.2 g/mL in lysis buffer to be flash frozen.

Resuspended membranes were thawed and homogenized using a dounce homogenizer, and n-Dodecyl-beta-Maltoside (DDM) (Anatrace, Cat#D310) added to a final concentration of 1% (w/v). After 30 min. at 8°C with gentle agitation, insoluble material was pelleted by ultracentrifugation (100,000 rcf, 30 min., 4°C) and the decanted supernatant passed through a 0.22 um filter.

The filtered supernatant was applied to a 1 mL StrepTrap HP column (GE, Cat#28-9075-47) equilibrated with binding buffer (100 mM HEPES pH 7.5, 150 mM NaCl, 0.03% DDM) and bound material eluted using elution buffer (100 mM HEPES pH 7.5, 150 mM NaCl, 0.03% DDM, 10 mM desthiobiotin (IBA, Cat#2-1000-002)). The peak elution fractions were pooled and incubated with HRV 3C protease (50 ug protease pr. 1 mg eluted protein) for 3 hrs at 8°C to cleave off the recombinant tag. The expression plasmid for HRV 3C protease (featuring an N-terminal 6xHis-tag) and protocols for expression and purification was kindly supplied by Jonathan Elegheert (then at University of Oxford).

#### Nanodisc reconstitution

To generate lipid nanodiscs, the HRV 3C digest was incubated with lipids and purified membrane scaffold protein (MSP1D1). Lipid stocks were prepared at ∼10 mM (assuming average M_w_ = 750 Da for individual lipid molecules) by drying 10 mg lipids (either Brain Polar Lipid Extract (Avanti Polar Lipids, Cat#141101) for ‘native’ nanodiscs or synthetic POPC (Avanti Polar Lipids, Cat#850457) for ‘minimal’ nanodiscs) suspended in chloroform under N_2_, and resuspending the dry pellet in 200 uL 10% DDM, 660 uL 2x TS buffer (20 mM Tris pH 7.5, 150 mM NaCl), and 470 uL MQ to a final volume of 1330 uL. For the BPL stocks, the 10% DDM stock was supplemented with 2% CHS (Anatrace, Cat#CH210), which resulted in a final lipid:CHS molar ratio of ∼60:40, which closely resembles the native lipid:cholesterol ratio in animal cell plasma membranes^74^. Lipids were resuspended by sonication. The membrane scaffold protein, MSP1D1, was expressed and purified according to the protocol published by Bayburt et. al^75^.

HRV 3C digested WT hMRP4, purified MSP1D1 and resuspended lipids were mixed in a 1:10:250 molar ratio: Protein and lipids were incubated for 1 hr at 8°C with gentle agitation, after which MSP1D1 was added. After incubation for 5 min at 8°C with gentle agitation, 50 mg (wet weight) freshly prepared SM2-resin Bio-Beads (BioRad, Cat#1523920) were added to adsorb detergent, and the suspension incubated for 16 hrs at 8°C with gentle agitation.

After incubation, the Bio-Beads were allowed to settle, and the suspension removed and passed once through a 30 µm cut off disposable Micro Bio-Spin filter (Bio-Rad, Cat#7326204) to get rid of any remaining beads. The filtered sample was injected onto a Superose 6 Increase 10/300 GL gel filtration column (GE Healthcare) equilibrated in nanodisc buffer (20 mM HEPES pH 7.5, 150 mM NaCl) and fractions containing complete nanodiscs (Fig. S1) pooled and concentrated using a Vivaspin 500 PES 50 kDa cut off spin filter (Sartorius, Cat#VS0131).

#### ATPase activity assays

Two assays were used for detection of ATP hydrolysis: The ADP-Glo assay (Promega, Cat#V6930), which detects ADP, and the PiColorLock assay (abcam, Cat#ab270004), which detects orthophosphate (see Fig. 2).

MTX interfered with the ADP-Glo assay (high luminescence counts were recorded even for the buffer controls). Consequently, the PiColorLock assay was used for samples containing MTX.

For both assays, three separate rounds of experiments were conducted (each with three technical replicates). Purified, nanodisc-reconstituted hMRP4 was resuspended in reaction buffer (20 mM Tris pH 7.5, 100 mM NaCl, 20 mM MgCl_2_, 20 mM KCl) at 130 nM, and 100 uM PGE2 (Selleck Chemicals, Cat#S3003), 100 uM TPT (Cayman Chemical, Cat#14129-50) or 1 mM MTX (Bio Basic Inc, Cat# MB0612) was added where relevant and the sample preheated at 37°C for 2 min. To start the reaction, 100 uM nucleotides (either UltraPure ATP (from the ADP-Glo kit) or AMP (Sigma, Cat#A1752)) were added, and the reaction allowed to proceed for 30 min. For each assayed condition, equivalent samples featuring no protein (buffer controls) were included to allow for subtraction of background signal.

To quench the ADP-Glo experiments, ADP-Glo Reagent was added, followed by ADP-Glo Detection Reagent, and luminescence counts (proportional to the amount of ADP produced during the reaction) were recorded using a Fluoroskan Microplate Fluorometer.

To quench the PiColorLock experiments, freshly prepared PiColorLock Reagent Mix was added, followed by addition of stabilizer. Development of color (proportional to the amount of orthophosphate produced during the reaction) was followed by measuring A635nm using a Fluoroskan Microplate Fluorometer.

#### Drug susceptibility assay

All cell lines used for this assay were based on the parental HEK293T LLP-Int-Blast cell line^76^ featuring a synthetic landing pad at the AAVS1 locus. This landing pad encodes a Tet-promoter preceding a Bxb1 attP recombination sequence, followed by the Bxb1 recombinase gene linked with the BFP gene and a blasticidin resistance gene (*Bsd*) via parechovirus 2A-like translational stop-start sequences. The Tet-promoter allows for doxycycline inducible expression of Bxb1 recombinase (as well as BFP and *Bsd*) until it facilitates integration of a transfected Bxb1 attB-containing donor plasmid and thus terminates its own expression. The integrity of the landing pad is ensured by continuous blasticidin resistance selection.

To generate derivative cell lines (featuring doxycycline inducible overexpression of hMRP4 constructs), HEK293T LLP-Int-Blast cells were transfected with a modified version of the attB-PuroR-2A-mCherry donor plasmid^76^ in which the mCherry cassette has been replaced with the hMRP4 cDNA (encoding either wildtype or mutant protein) in tandem with an mVenus-tag linked in tandem to a Twin-Strep-tag and a 6xHis-tag. The fluorescent tag allows for verification of expression using fluorescence microscopy, while the Twin-Strep-tag allows for assessment of expression levels using Western blot.

Doxycycline dependent integration of the transfected donor plasmid (featuring the hMRP4 construct and the 2A-linked puromycin resistance gene) to generate the derivative HEK293T LLP-Puro-MRP4 cell line is verified by fluorescence microscopy and puromycin resistance selection.

To conduct the drug susceptibility assays, HEK293T LLP-Puro-MRP4 (encoding either wildtype or mutant recombinant hMRP4) and HEK293T LLP-Int-Blast (the parental cell line acting as negative control, NEG) were seeded in poly-L-ornithine (Sigma, Cat#P3655) coated 96-well plates at 40,000 cells/well (100 uL volume) and incubated at 37°C for 24 hrs. Stocks of MTX (ranging from 0.005 – 0.5 mM) were generated from a 100 mM stock (in DMSO), 10x diluted (and pH-adjusted to pH 7.5) in PBS, and appropriately diluted in plating medium immediately prior to use (final [DMSO] < 0.5%).

To initiate MTX challenge, incubation medium was replaced with MTX-containing medium (100 uL volume) and the plates incubated at 37°C for 4 hrs. To terminate MTX challenge, MTX-containing medium was removed, and each well washed with 2×100 uL PBS before fresh plating medium (100 uL volume) was added and the plates left at 37°C for 44 hrs.

Cell Proliferation Kit I (MTT) (Roche, Cat#11465007001) was used to assess cell viability 48 hrs post initiation of MTX challenge: Incubation medium was removed and replaced with plating medium (100 uL volume) supplemented with 0.5 mg/mL MTT and the plates left at 37°C to allow for conversion of MTT into Formazan. After 2 hrs, 100 uL detergent solution was added to each well and the plates left at 37°C for 22 hrs. To quantify cell viability, the A600nm of each well was recorded using a Fluoroskan Microplate Fluorometer.

#### Electron microscopy sample preparation

All the samples used for cryo-EM were prepared using freshly purified and concentrated nanodisc-reconstituted hMRP4 and freshly prepared stocks of ATP (Bio Basic Inc, Cat#AB0020), ATPγS (Roche, Cat#11162306001), MgCl_2_, PGE2, TPT and MTX in stock buffer (100 mM HEPES pH 7.5, 150 mM NaCl).

For the sample used to obtain the reconstruction of IF MRP4 (no substrate added) in native nanodiscs (see Fig. S2, S8A, and S9D), WT hMRP4 in native nanodiscs was incubated with 10 mM ATPγS and 10 mM MgCl_2_ for 30 min on ice prior to grid preparation.

For the sample used to obtain the (low resolution) reconstruction of IF MRP4 in minimal nanodiscs (nothing added) (see Fig. S8B), WT hMRP4 in minimal nanodiscs was incubated for 30 min on ice prior to grid preparation.

For the sample used to obtain the additional reconstruction of IF MRP4 (nothing added) (see Fig. S8C and S9E), a separate preparation of WT hMRP4 in native nanodiscs was incubated for 30 min on ice prior to grid preparation.

For the sample used to determine the structure of OF MRP4, E1202Q hMRP4 in minimal nanodiscs was incubated with 10 mM ATP, 10 mM MgCl_2_, and 100 uM TPT for 5 min at 37°C (to mimic turnover conditions) prior to grid preparation.

For the samples used to determine the structures of IF MRP4 (MTX or TPT added), WT hMRP4 in minimal nanodiscs was incubated with 10 mM ATP, 10 mM MgCl_2_, and either 100 uM MTX or 100 uM TPT for 5 min at 37°C prior to grid preparation. For the sample used to determine the structure of IF MRP4 (PGE2 added), WT hMRP4 in minimal nanodiscs was incubated with 1 mM PGE2 for 5 min at 37°C prior to grid preparation.

In all cases, sample was applied to a freshly glow discharged R2/1 Cu 300 Quantifoil grid (Plano GmbH, Cat#S174-2), and a Vitrobot Mark IV (FEI) was used to blot away excess sample and immediately plunge the grids into liquid ethane for vitrification of embedded sample.

#### Cryo-EM data collection and processing

Movies were acquired using a Titan Krios G2 (FEI) fitted with a Falcon 3EC detector operated in counting mode. Data acquisition was semi-automated using EPU (FEI, Thermo Scientific). Relevant details for the different datasets are summarized in Table S1 and S2.

The final reconstructions were obtained using cryoSPARC^77^ as described in Fig. S2-6. Global and directional resolution of the final Coulomb potential maps was assessed using the Remote 3DFSC Processing Server^78^ (https://3dfsc.salk.edu/). Local resolution of the final maps was calculated using cryoSPARC (using an FSC threshold of 0.143) and visualized using UCSF Chimera^79^. Euler angle distribution plots were generated using cryoSPARC.

#### Model building and refinement

MODELLER^80^ was used to generate a homology model of IF hMRP4 based on IF bMRP1 (PDB: 5UJ9), which was used as a starting point for model building of all four IF MRP4 atomic models. Likewise, a homology model of OF hMRP4 based on OF bMRP1 (PDB: 6BHU) was generated and used as a starting point for model building of the OF MRP4 structure. The homology models were rigid-body fit into the respective maps using UCSF Chimera^79^ and real-space refined in PHENIX^81^. The models were then inspected visually and manually fitted in Coot^82^.

For further optimized fitting of the IF MRP4 structures, the online molecular dynamics pipeline, Namdinator^83^, was used. In all cases, the NBD1 moiety of the models could not be fitted reliably into the maps using this method. Consequently, NBD1 of the IF hMRP4 models was deleted and the remainder of the models fitted into the respective maps in UCSF Chimera. This was followed by fitting the respective NBD1 domains separately, while taking care of bond continuity for each of the complete models. Further fitting of amino acid side chains was done in iterative cycles of manual model building into the map using Coot and ISOLDE^84^.

All ligands were fitted into map density in Coot and restraints generated in eLBOW (PHENIX), followed by real-space refinement of the complete models in PHENIX. MolProbity^85^ was used to assess the quality of the final models. Maps and models were visualized using UCSF Chimera. The DelPhi Web Server^86^ (http://compbio.clemson.edu/sapp/delphi_webserver/) was used for calculating the electrostatic energies (pH = 7.0, salt concentration = 0.15 M, default dielectric constant values) displayed in the electrostatic surface representation of IF hMRP4 (no substrate added) (visualized using UCSF Chimera).

#### Figure preparation

Figures were prepared using UCSF Chimera^79^, GraphPad Prism 9 (GraphPad Software), and Illustrator (Adobe). Elements of Fig. 4 were obtained from BioRender.com.

### QUANTIFICATION AND STATISTICAL ANALYSIS

For the ATPase activity assays, the data points represent the average values obtained from independent experiments (each with three technical replicates). Three rounds of independent experiments were performed for each condition.

For the drug susceptibility assay, three separate estimates of pIC_50_ for each cell line were obtained from three rounds of independent experiments (each with six technical replicates), with each estimate obtained from fitting a non-linear sigmoidal dose-response model (using GraphPad Prism 9) to the data points.

In order to phenotype the cell lines, a Šídák’s multiple comparisons test was performed (using GraphPad Prism 9), comparing selected pairs of mean pIC_50_ values: The mean pIC_50_ values (the average of three independent estimates obtained from the fits displayed in Fig. S10) of the individual cell lines were simultaneously compared to the mean pIC_50_ values determined for the wildtype (WT) and parental (NEG) cell lines, that were chosen to represent the defining pIC_50_ for, respectively, a resistant (R phenotype) cell line and a sensitive (S phenotype) cell line.

The results of the Šídák’s multiple comparisons test are thus interpreted:

If the mean pIC_50_ of a cell line is not significantly different from the mean pIC_50_ of the WT cell line, but at the same time is significantly different from the mean pIC_50_ of the NEG cell line, the cell line is said to display an R phenotype.

If the mean pIC_50_ of a cell line is significantly different from the mean pIC_50_ of the WT cell line, but at the same time is not significantly different from the mean pIC_50_ of the NEG cell line, the cell line is said to display an S phenotype.

If the mean pIC_50_ of a cell line is not significantly different from the mean pIC_50_ of the WT cell line, and also is not significantly different from the mean pIC_50_ of the NEG cell line, the phenotyping is said to be inconclusive (X).

