## Supplemental information for "Structural and Mechanistic Basis of Substrate Transport by the Multidrug Transporter MRP4"

SUPPLEMENTAL FIGURES (TITLES AND LEGENDS)

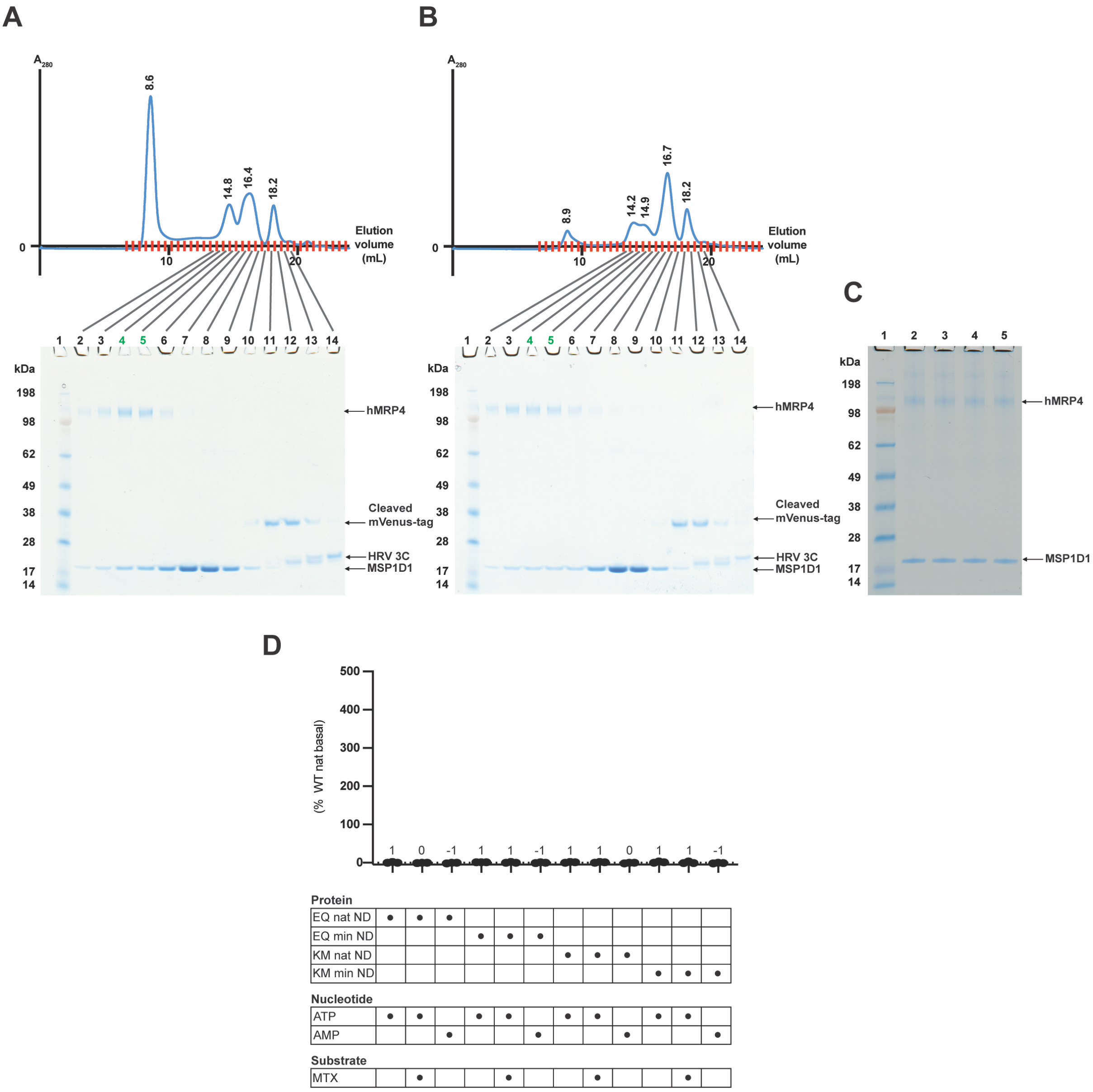

**Fig. S1: Preparation of nanodisc-reconstituted hMRP4.**
Analysis of reconstitution of WT hMRP4 in (A) native and (B) minimal nanodiscs.
Top: Typical gel filtration chromatograms. The elution volume corresponding to the maximum $A_{280}$ for each separate peak is indicated above it, and the elution fractions are marked with red ticks.
Bottom: SDS PAGE analysis of individual elution fractions. For both reconstitutions, typically, the elution fractions analyzed in lanes 4 and 5 of the respective SDS PAGE gels were pooled and concentrated for cryo-EM sample preparation and ATPase activity assays. The molecular identity of the individual bands are indicated.
$M_w$(hMRP4) = 150 kDa, $M_w$(Cleaved mVenus-tag) = 31 kDa, $M_w$(HRV 3C protease) = 24 kDa, $M_w$(MSP1D1) = 22 kDa.
(C) SDS PAGE analysis of representative concentrated samples of nanodisc-reconstituted hMRP4. Lane 1: Molecular weight marker. Lane 2: E1202Q (EQ) hMRP4 in native nanodiscs. Lane 3: EQ hMRP4 in minimal nanodiscs. Lane 4:

K1081M (KM) hMRP4 in native nanodiscs. Lane 5: KM hMRP4 in minimal nanodiscs.

(D) Basal and substrate-modulated ATPase activity of the EQ and KM variants of hMRP4 assayed using the PiColorLock assay. Data are presented as described in Fig. 2. Experiments were performed as described in Fig. 2 and in parallel with the experiments presented in Fig. 2.

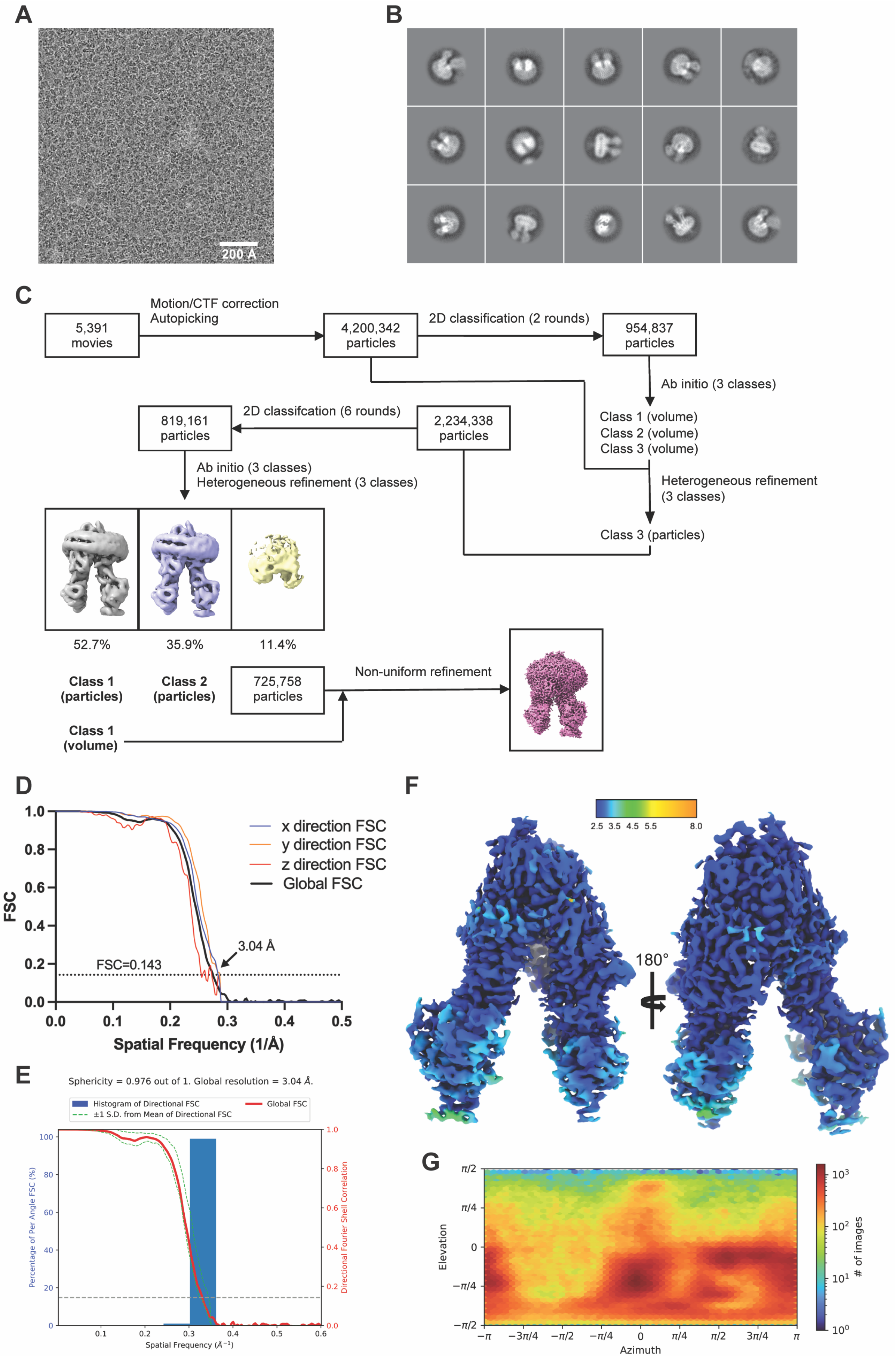

A
200 Å
B
C
5,391 movies
Motion/CTF correction
Autopicking
4,200,342 particles
2D classification (2 rounds)
954,837 particles
Ab initio (3 classes)
Class 1 (volume)
Class 2 (volume)
Class 3 (volume)
Heterogeneous refinement (3 classes)
Class 3 (particles)
2,234,338 particles
2D classifcation (6 rounds)
819,161 particles
Ab initio (3 classes)
Heterogeneous refinement (3 classes)
52.7%
35.9%
11.4%
Class 1 (particles)
Class 2 (particles)
725,758 particles
Non-uniform refinement
Class 1 (volume)
D
FSC
x direction FSC
y direction FSC
z direction FSC
Global FSC
3.04 Å
FSC=0.143
Spatial Frequency (1/Å)
E
Sphericity = 0.976 out of 1. Global resolution = 3.04 Å.
Histogram of Directional FSC
±1 S.D. from Mean of Directional FSC
Global FSC
Percentage of Per Angle FSC (%)
Directional Fourier Shell Correlation
Spatial Frequency (Å⁻¹)
F
2.5 3.5 4.5 5.5 8.0
180°
G
Elevation
Azimuth
### of images

**Fig. S2: Overview of the data collection and processing that resulted in the final reconstruction of IF MRP4 (no substrate added).**
(A) Representative micrograph (with scale bar). (B) Representative 2D classes. (C) Overview of the data processing pipeline in cryoSPARC featuring images of intermediate and final maps. (D) Global and directional Fourier shell correlation (FSC) curves for the final map. The global resolution (as calculated using an FSC threshold of 0.143 (punctured line)) is stated and indicated (arrow). (E) Plot of the global FSC curve (red line) including the ±1 standard deviation spread of directional resolution (green punctured lines) and a histogram (blue) of 100 evenly sampled directional resolutions. (F) The final Coulomb potential map colored by local resolution (in Å). (G) Euler angle distribution plot.

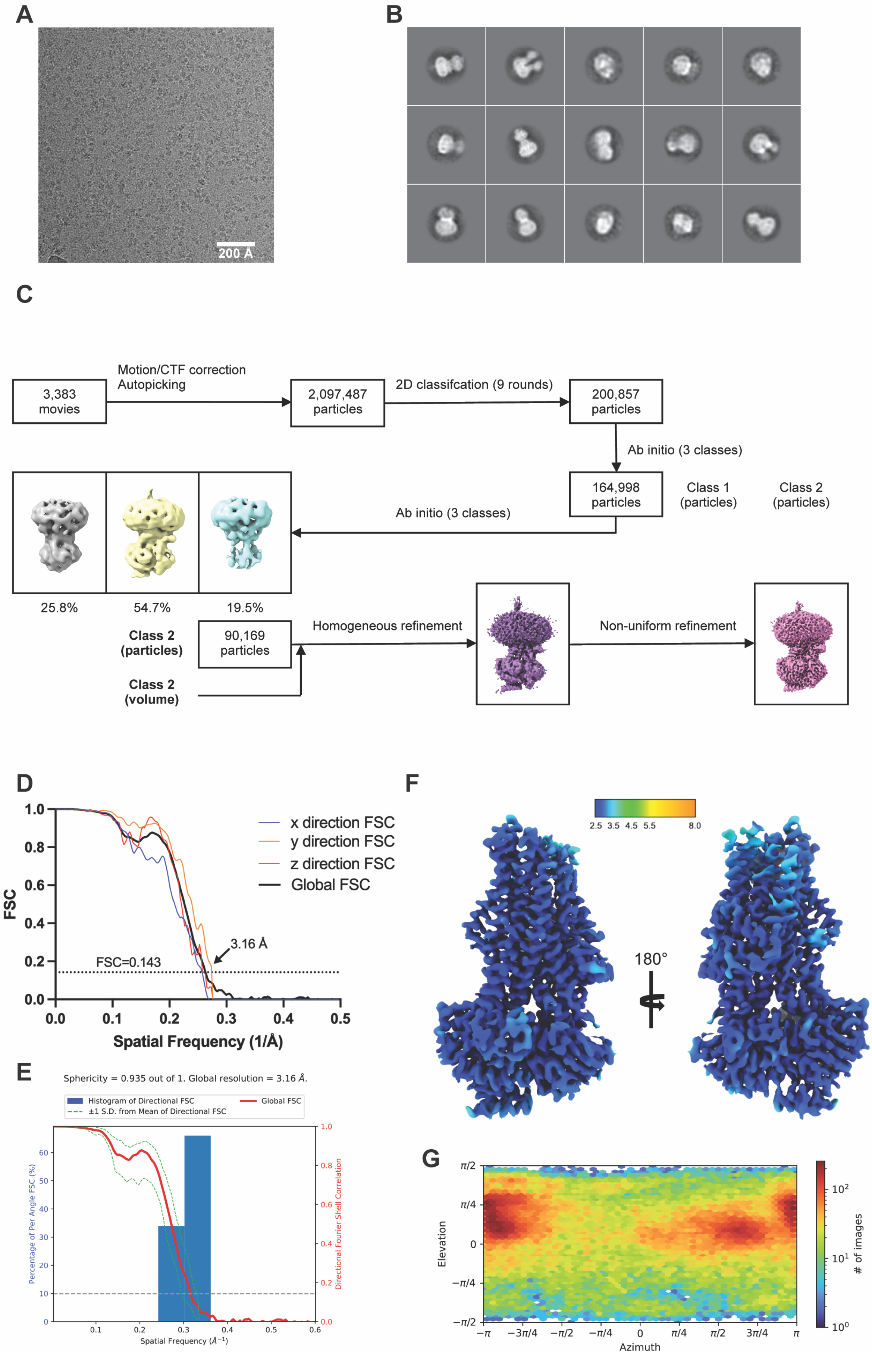

A
200 Å
B
C
3,383 movies
Motion/CTF correction
Autopicking
2,097,487 particles
2D classifcation (9 rounds)
200,857 particles
Ab initio (3 classes)
164,998 particles
Class 1 (particles)
Class 2 (particles)
Ab initio (3 classes)
25.8%
54.7%
19.5%
Class 2 (particles)
90,169 particles
Class 2 (volume)
Homogeneous refinement
Non-uniform refinement
D
FSC
Spatial Frequency (1/Å)
x direction FSC
y direction FSC
z direction FSC
Global FSC
3.16 Å
FSC=0.143
E
Sphericity = 0.935 out of 1. Global resolution = 3.16 Å.
Histogram of Directional FSC
±1 S.D. from Mean of Directional FSC
Global FSC
Percentage of Per Angle FSC (%)
Directional Fourier Shell Correlation
Spatial Frequency (Å⁻¹)
F
2.5 3.5 4.5 5.5 8.0
180°
G
Elevation
Azimuth
### of images

**Fig. S3: Overview of the data collection and processing that resulted in the final reconstruction of OF MRP4.**

(A) Representative micrograph (with scale bar). (B) Representative 2D classes. (C) Overview of the data processing pipeline in cryoSPARC featuring images of intermediate and final maps. (D) Global and directional Fourier shell correlation (FSC) curves for the final map. The global resolution (as calculated using an FSC threshold of 0.143 (punctured line)) is stated and indicated (arrow). (E) Plot of the global FSC curve (red line) including the ±1 standard deviation spread of directional resolution (green punctured lines) and a histogram (blue) of 100 evenly sampled directional resolutions. (F) The final Coulomb potential map colored by local resolution (in Å). (G) Euler angle distribution plot.

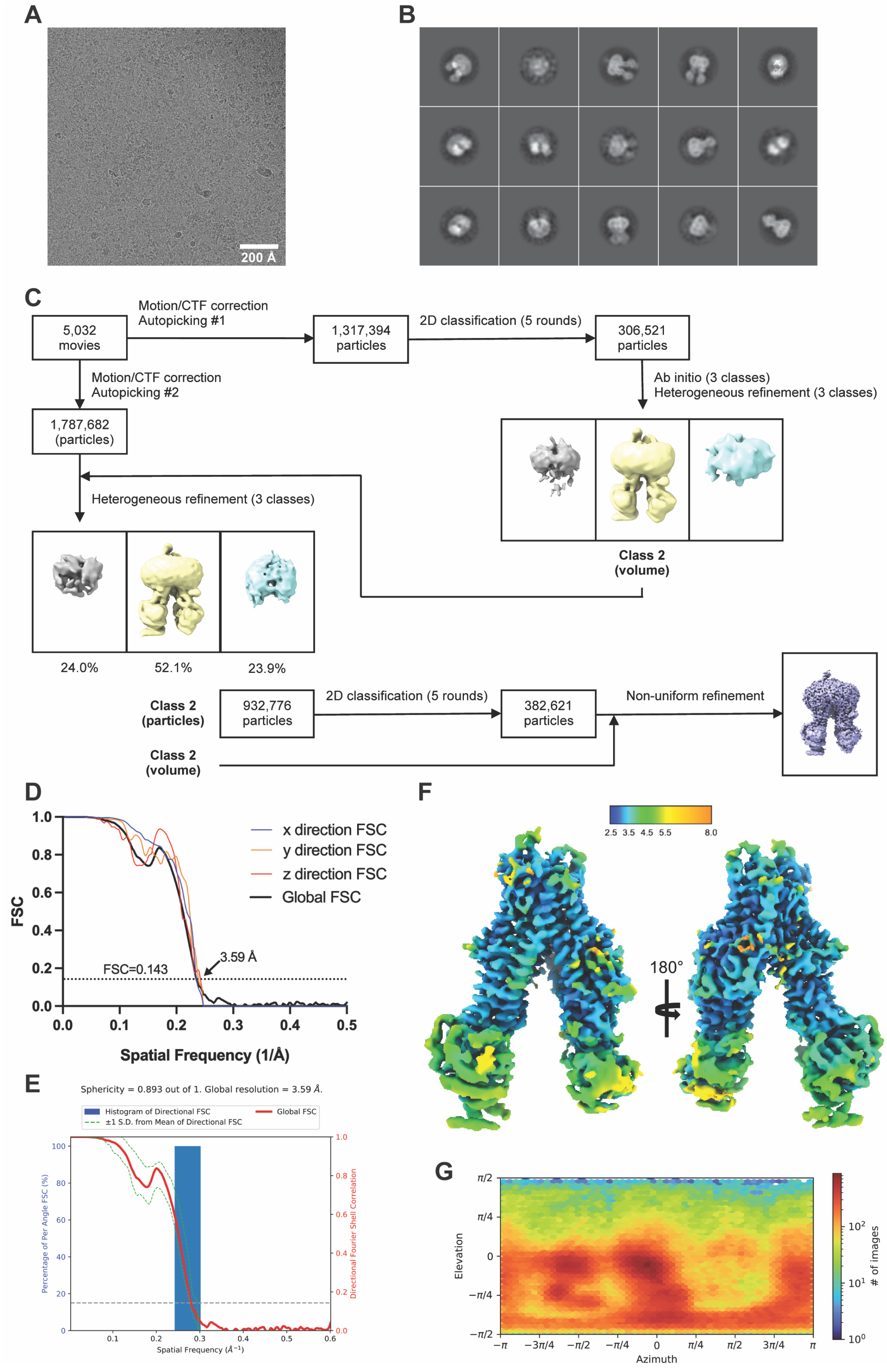

A
200 Å
B
C
5,032 movies
Motion/CTF correction
Autopicking #1
1,317,394 particles
2D classification (5 rounds)
306,521 particles
Motion/CTF correction
Autopicking #2
1,787,682 (particles)
Ab initio (3 classes)
Heterogeneous refinement (3 classes)
Heterogeneous refinement (3 classes)
Class 2 (volume)
24.0%
52.1%
23.9%
Class 2 (particles)
932,776 particles
2D classification (5 rounds)
382,621 particles
Non-uniform refinement
Class 2 (volume)
D
FSC
x direction FSC
y direction FSC
z direction FSC
Global FSC
3.59 Å
FSC=0.143
Spatial Frequency (1/Å)
E
Sphericity = 0.893 out of 1. Global resolution = 3.59 Å.
Histogram of Directional FSC
±1 S.D. from Mean of Directional FSC
Global FSC
Percentage of Per Angle FSC (%)
Directional Fourier Shell Correlation
Spatial Frequency (Å⁻¹)
F
2.5 3.5 4.5 5.5 8.0
180°
G
Elevation
Azimuth
### of images

**Fig. S4: Overview of the data collection and processing that resulted in the final reconstruction of IF MRP4 (MTX added).**
(A) Representative micrograph (with scale bar). (B) Representative 2D classes. (C) Overview of the data processing pipeline in cryoSPARC featuring images of intermediate and final maps. (D) Global and directional Fourier shell correlation (FSC) curves for the final map. The global resolution (as calculated using an FSC threshold of 0.143 (punctured line)) is stated and indicated (arrow). (E) Plot of the global FSC curve (red line) including the ±1 standard deviation spread of directional resolution (green punctured lines) and a histogram (blue) of 100 evenly sampled directional resolutions. (F) The final Coulomb potential map colored by local resolution (in Å). (G) Euler angle distribution plot.

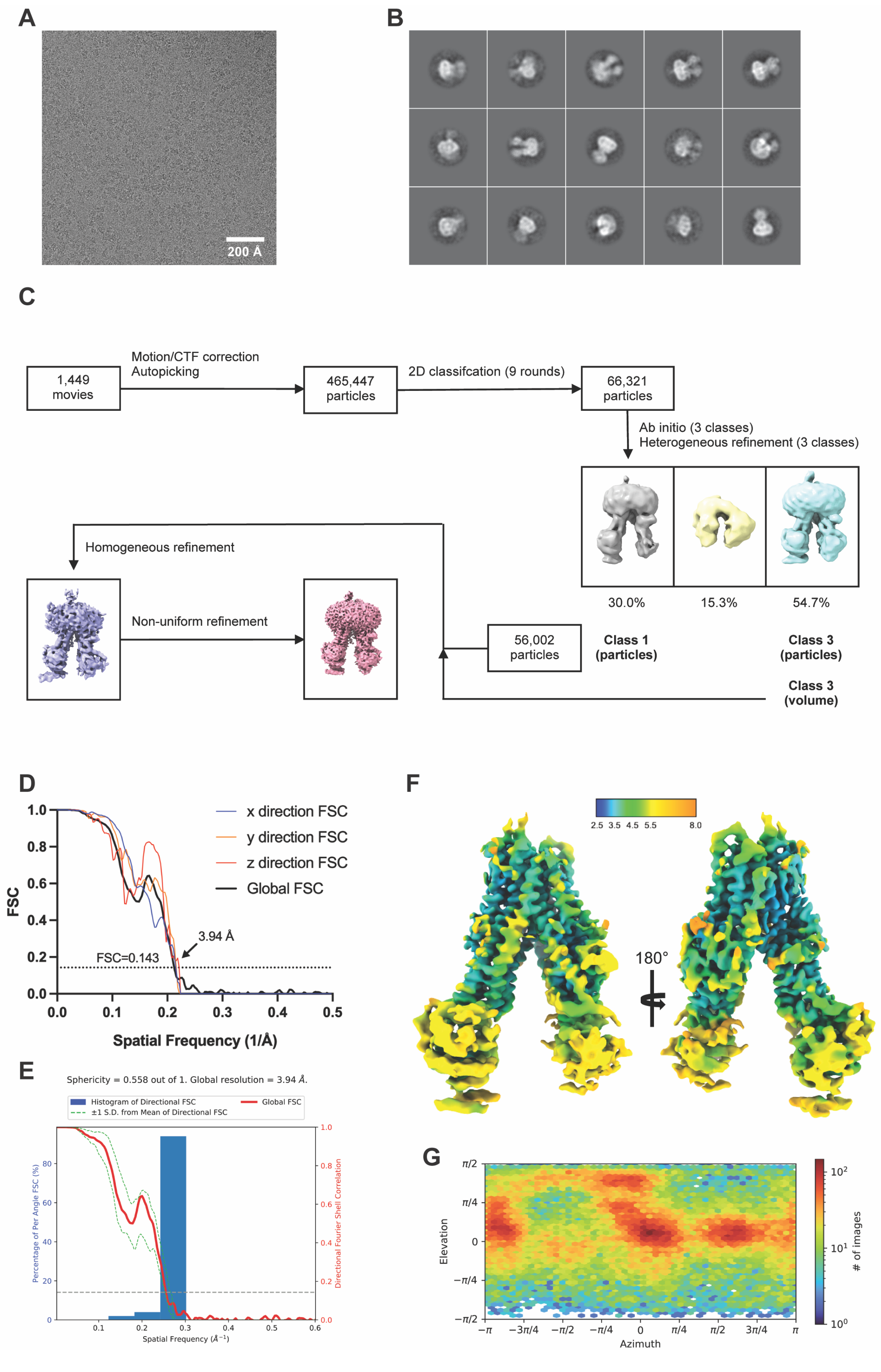
A
200 Å
B
C
1,449 movies
Motion/CTF correction
Autopicking
465,447 particles
2D classifcation (9 rounds)
66,321 particles
Ab initio (3 classes)
Heterogeneous refinement (3 classes)
30.0%
15.3%
54.7%
Homogeneous refinement
Non-uniform refinement
56,002 particles
Class 1 (particles)
Class 3 (particles)
Class 3 (volume)
D
FSC
x direction FSC
y direction FSC
z direction FSC
Global FSC
3.94 Å
FSC=0.143
Spatial Frequency (1/Å)
E
Sphericity = 0.558 out of 1. Global resolution = 3.94 Å.
Histogram of Directional FSC
±1 S.D. from Mean of Directional FSC
Global FSC
Percentage of Per Angle FSC (%)
Directional Fourier Shell Correlation
Spatial Frequency (Å⁻¹)
F
2.5 3.5 4.5 5.5 8.0
180°
G
Elevation
Azimuth
### of images

**Fig. S5: Overview of the data collection and processing that resulted in the final reconstruction of IF MRP4 (TPT added).**
(A) Representative micrograph (with scale bar). (B) Representative 2D classes. (C) Overview of the data processing pipeline in cryoSPARC featuring images of intermediate and final maps. (D) Global and directional Fourier shell correlation (FSC) curves for the final map. The global resolution (as calculated using an FSC threshold of 0.143 (punctured line)) is stated and indicated (arrow). (E) Plot of the global FSC curve (red line) including the ±1 standard deviation spread of directional resolution (green punctured lines) and a histogram (blue) of 100 evenly sampled directional resolutions. (F) The final Coulomb potential map colored by local resolution (in Å). (G) Euler angle distribution plot.

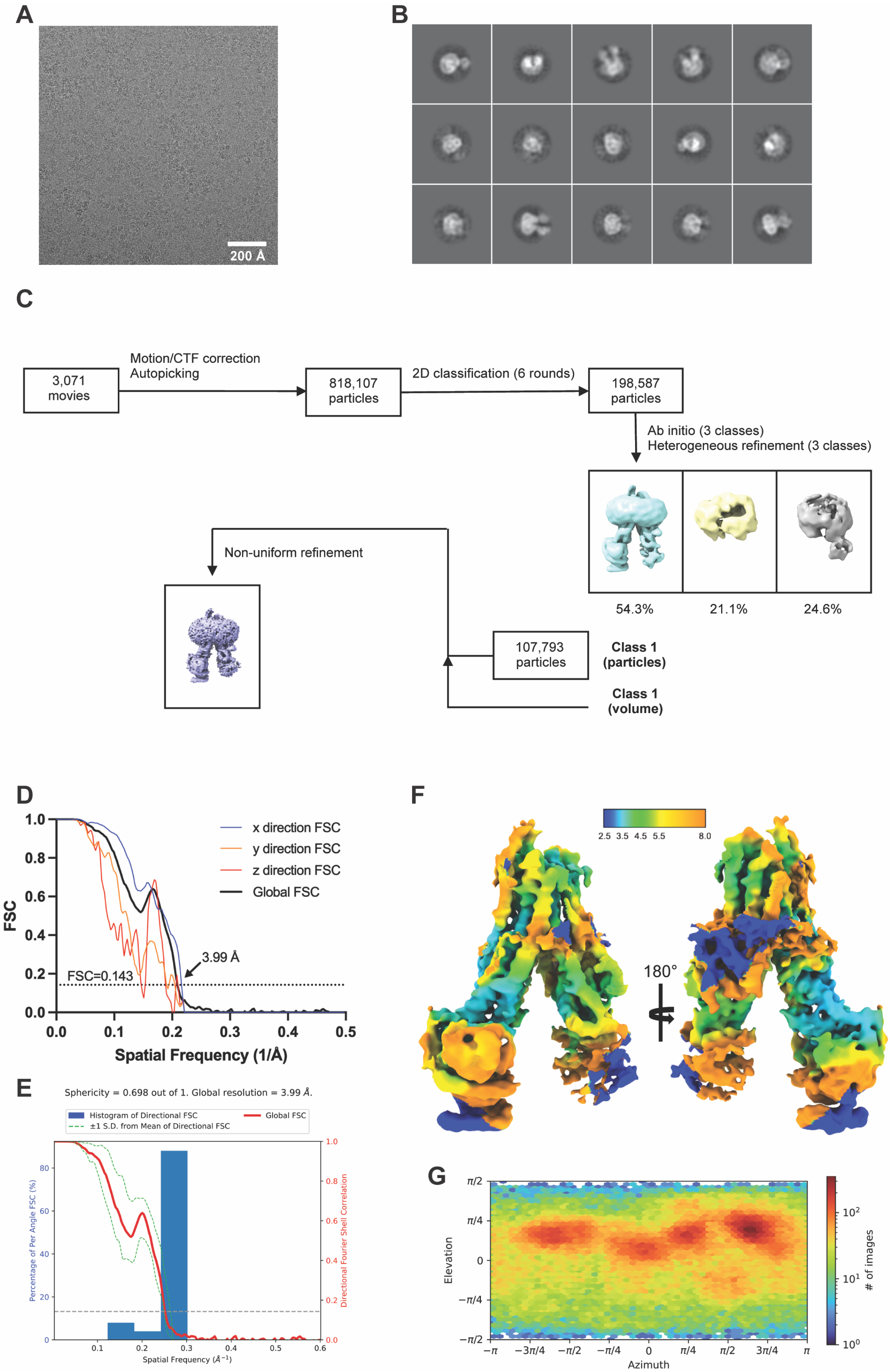
A
200 Å
B
C
3,071
movies
Motion/CTF correction
Autopicking
818,107
particles
2D classification (6 rounds)
198,587
particles
Ab initio (3 classes)
Heterogeneous refinement (3 classes)
54.3%
21.1%
24.6%
Non-uniform refinement
107,793
particles
Class 1
(particles)
Class 1
(volume)
D
FSC
x direction FSC
y direction FSC
z direction FSC
Global FSC
3.99 Å
FSC=0.143
Spatial Frequency (1/Å)
E
Sphericity = 0.698 out of 1. Global resolution = 3.99 Å.
Histogram of Directional FSC
±1 S.D. from Mean of Directional FSC
Global FSC
Percentage of Per Angle FSC (%)
Directional Fourier Shell Correlation
Spatial Frequency (Å⁻¹)
F
2.5 3.5 4.5 5.5 8.0
180°
G
Elevation
Azimuth
### of images

**Fig. S6: Overview of the data collection and processing that resulted in the final reconstruction of IF MRP4 (PGE2 added).**
(A) Representative micrograph (with scale bar). (B) Representative 2D classes. (C) Overview of the data processing pipeline in cryoSPARC featuring images of intermediate and final maps. (D) Global and directional Fourier shell correlation (FSC) curves for the final map. The global resolution (as calculated using an FSC threshold of 0.143 (punctured line)) is stated and indicated (arrow). (E) Plot of the global FSC curve (red line) including the ±1 standard deviation spread of directional resolution (green punctured lines) and a histogram (blue) of 100 evenly sampled directional resolutions. (F) The final Coulomb potential map colored by local resolution (in Å). (G) Euler angle distribution plot.

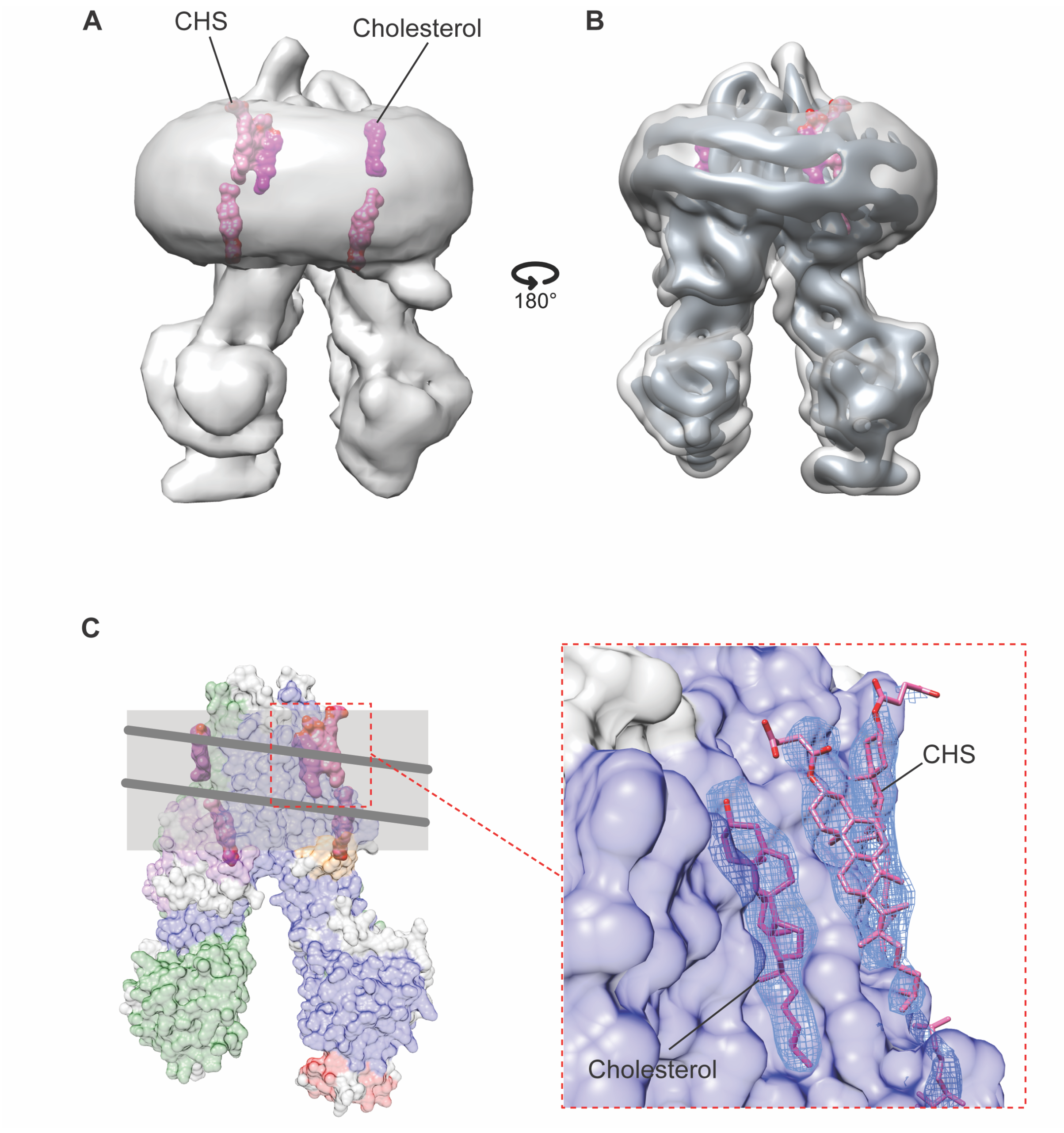

**Fig. S7: Structural characterization of nanodisc scaffold/bilayer and annular lipids.**
Various representations of the final reconstruction (Fig. S2) obtained from a sample of WT hMRP4 in native nanodiscs (no substrate added).
(A) Low-pass filtered (>20 Å) Coulomb potential map (contoured at $\sigma$ = 0.08) of IF MRP4 (no substrate added) featuring surface representations of modelled annular lipids (pink = CHS, magenta = cholesterol).
(B) Rotated view of (A) featuring an additional copy of the same low-pass filtered map (contoured at $\sigma$ = 0.18) displaying density corresponding to two copies of the MSP1D1 scaffold protein.
(C) Left: Surface representation of IF MRP4 (no substrate added) and annular lipids overlaid with outlines of lipid bilayer (light gray) and scaffold proteins (dark gray). Right: Zoomed in view of the outer bilayer leaflet with blue mesh representing map density (contoured at $\sigma$ = 0.50) corresponding to bound lipids (shown as sticks).

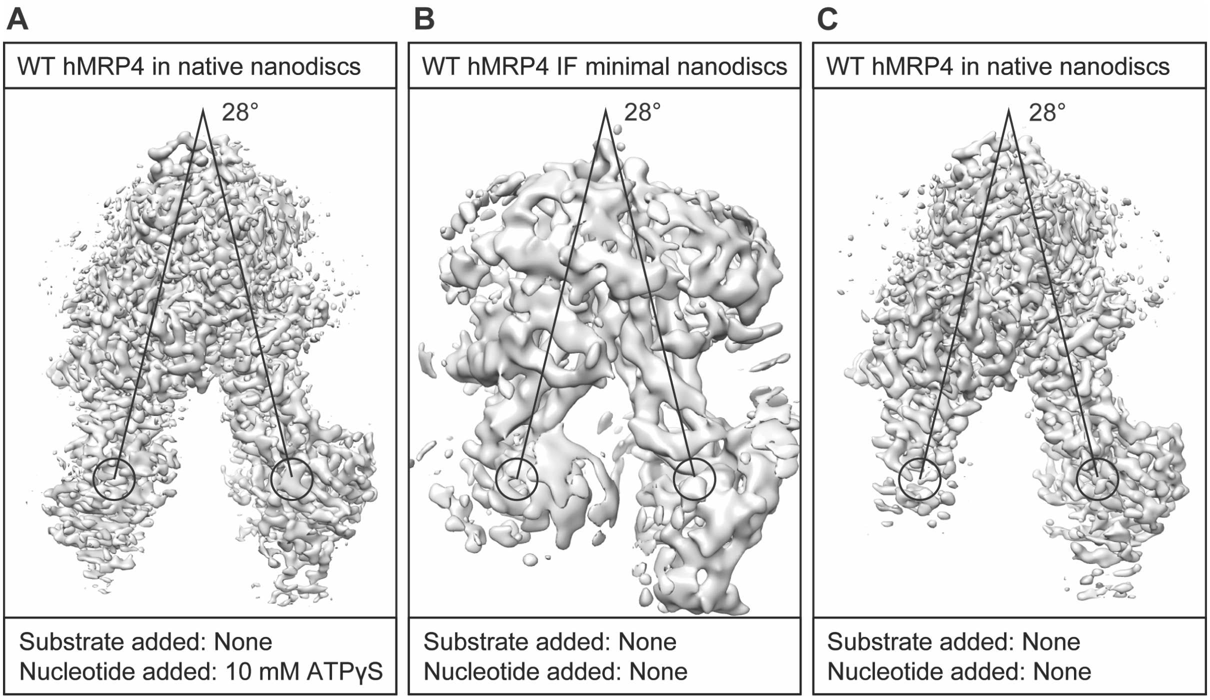

**Fig. S8: Reconstructions obtained from samples of WT hMRP4 reconstituted in native and minimal nanodiscs.**
(A) The final reconstruction (Fig. S2) obtained from a sample of WT hMRP4 reconstituted in native nanodiscs (contoured at $\sigma$ = 0.40). While no substrate was added to the sample, ATPγS and $MgCl_2$ was present at 10 mM (see Method details).
(B) The final reconstruction obtained from a sample of WT hMRP4 reconstituted in minimal nanodiscs (contoured at $\sigma$ = 0.40).
(C) The final reconstruction (also shown in Fig. S9E) obtained from a separate sample of WT hMRP4 reconstituted in native nanodiscs (contoured at $\sigma$ = 0.40).

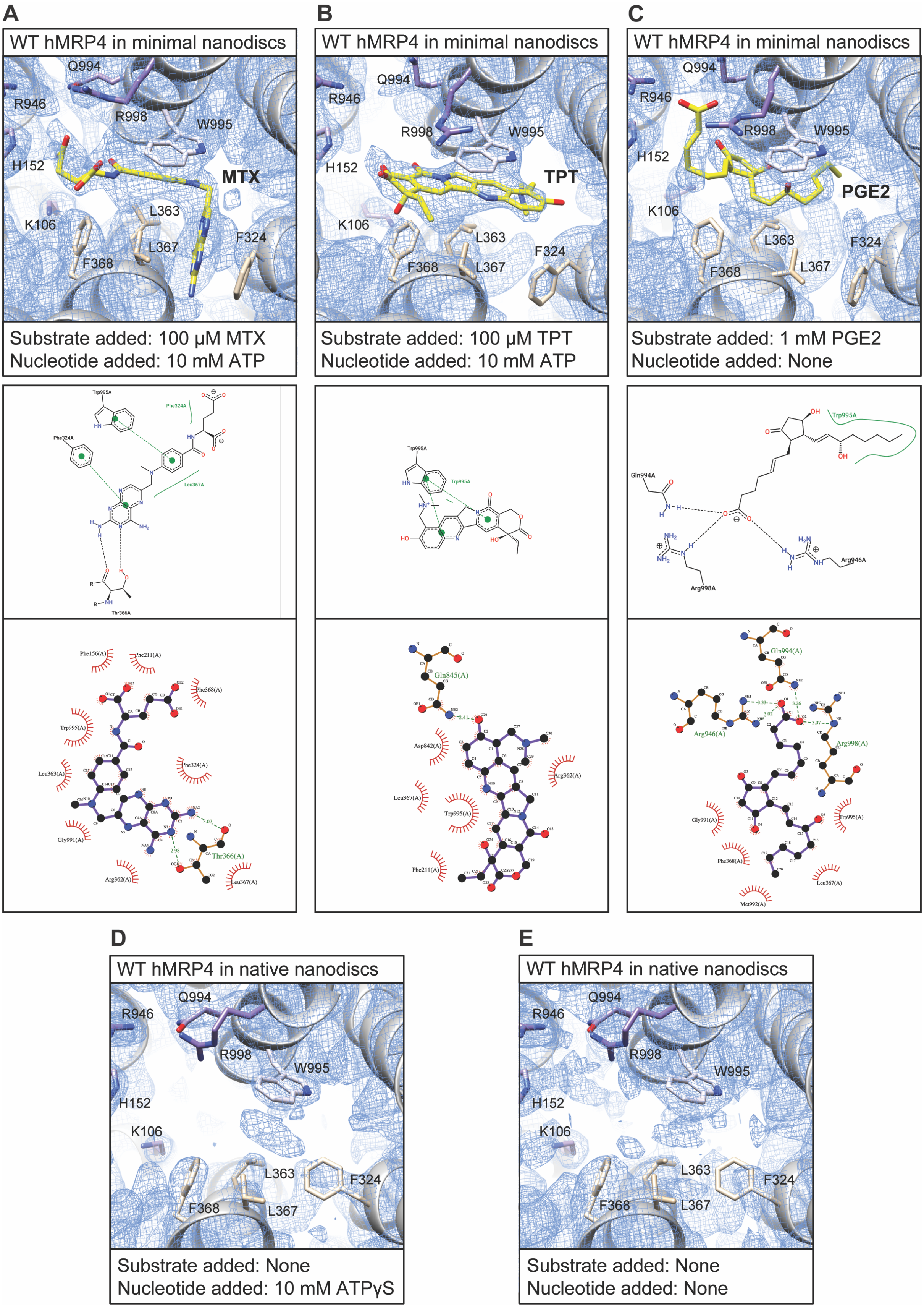
A
WT hMRP4 in minimal nanodiscs
Q994
R946
R998
W995
H152
MTX
K106
L363
L367
F324
F368
Substrate added: 100 μM MTX
Nucleotide added: 10 mM ATP
B
WT hMRP4 in minimal nanodiscs
Q994
R946
R998
W995
H152
TPT
K106
L363
F324
F368
L367
Substrate added: 100 μM TPT
Nucleotide added: 10 mM ATP
C
WT hMRP4 in minimal nanodiscs
Q994
R946
R998
W995
H152
PGE2
K106
L363
F324
F368
L367
Substrate added: 1 mM PGE2
Nucleotide added: None
D
WT hMRP4 in native nanodiscs
Q994
R946
R998
W995
H152
K106
L363
F324
F368
L367
Substrate added: None
Nucleotide added: 10 mM ATPγS
E
WT hMRP4 in native nanodiscs
Q994
R946
R998
W995
H152
K106
L363
F324
F368
L367
Substrate added: None
Nucleotide added: None

**Fig. S9: Characterization of the substrate binding pocket of obtained reconstructions of IF hMRP4.**
(A/B/C) The substrate binding pocket of the reconstructions (Fig. S4/S5/S6) obtained from samples of WT hMRP4 (reconstituted in minimal nanodiscs) to which MTX/TPT/PGE2 had been added (see Method details).
Top: Zoomed in view of maps (blue mesh, contoured at $\sigma$ = 0.40/0.27/0.20) and models (ribbons and sticks). 10 key residues of the substrate binding pocket are shown as sticks (colored according to Kyte-Doolittle hydrophobicity, see Fig. 3), and the fits of the respective substrate molecules (MTX/TPT/PGE2 (yellow sticks)) into the maps are shown.
Middle: PoseView[1] representation of the interaction between the MRP4 model and the modelled substrate molecule.
Bottom: LigPlot+[2] representation of the same interaction.
(D) The substrate binding pocket of the reconstruction (Fig. S2) obtained from a sample of WT hMRP4 (reconstituted in native nanodiscs). While no substrate was added to the sample, ATP$\gamma$S and $MgCl_2$ was present at 10 mM (see Method details). Zoomed in view of the Coulomb potential map (blue mesh, contoured at $\sigma$ = 0.80) and the corresponding model (ribbons and sticks) of IF MRP4 (no substrate added).
(E) The substrate binding pocket of a reconstruction (also shown in Fig. S8C) obtained from a separate sample of WT hMRP4 (reconstituted in native nanodiscs). Zoomed in view of the final Coulomb potential map (blue mesh, contoured at $\sigma$ = 0.35) overlaid with the model (ribbons and sticks) of IF MRP4 (no substrate added).

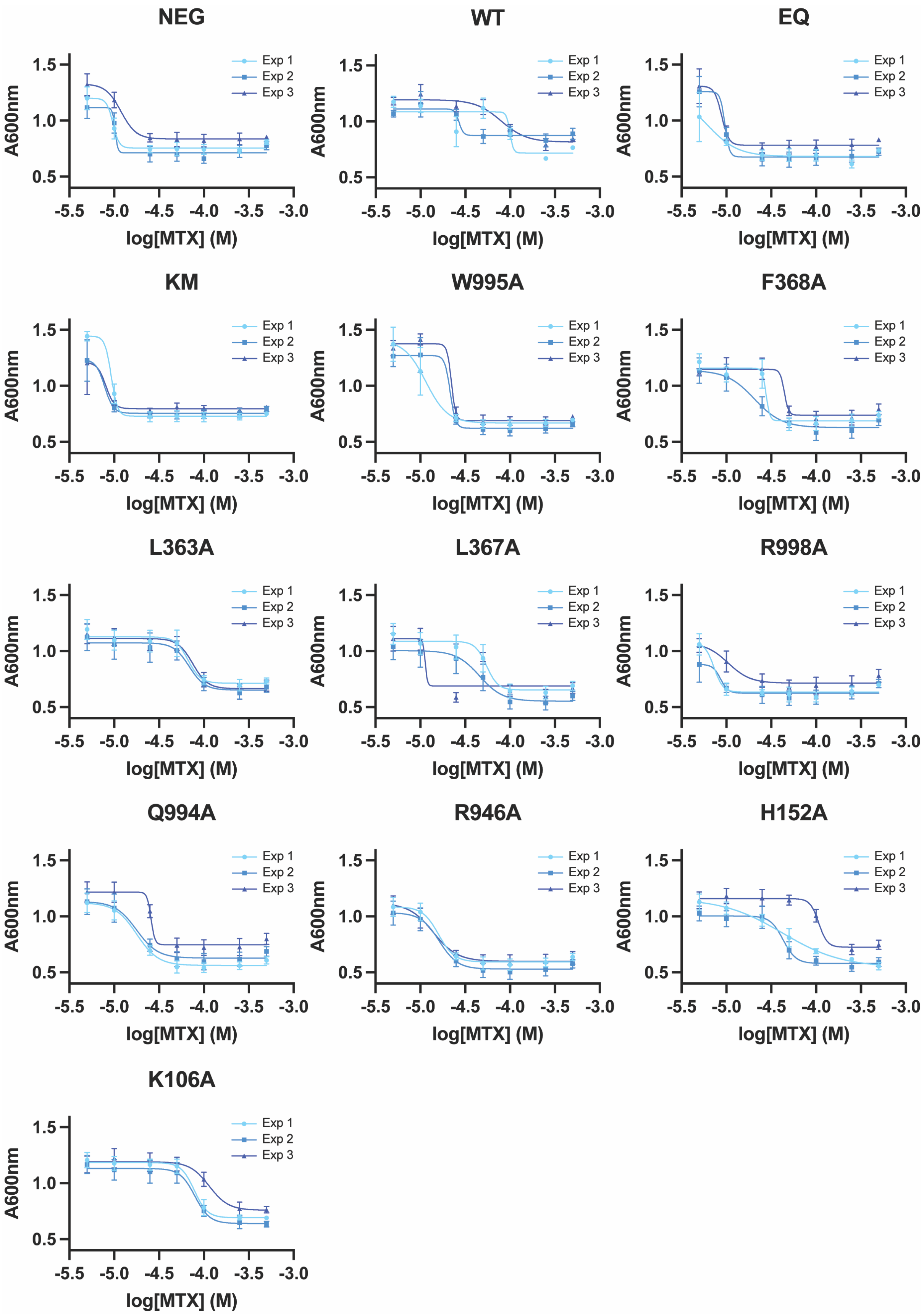
NEG
WT
EQ
KM
W995A
F368A
L363A
L367A
R998A
Q994A
R946A
H152A
K106A
Exp 1
Exp 2
Exp 3
A600nm
log[MTX] (M)
1.5
1.0
0.5
-5.5
-5.0
-4.5
-4.0
-3.5
-3.0

**Fig. S10: Plots of data and respective non-linear fits from which pIC$_{50}$ estimates were obtained.**
Plots of the data obtained from drug susceptibility assays of individual cell lines (identity indicated above plot) performed in parallel. Three separate rounds of experiments (Exp1-3) were performed for each cell line, with each data point representing an independent experiment (each with six technical replicates) and error bars representing the standard deviation. Non-linear sigmoidal dose-response curves were fitted to the data to obtain three separate estimates of pIC$_{50}$ for the respective cell line.

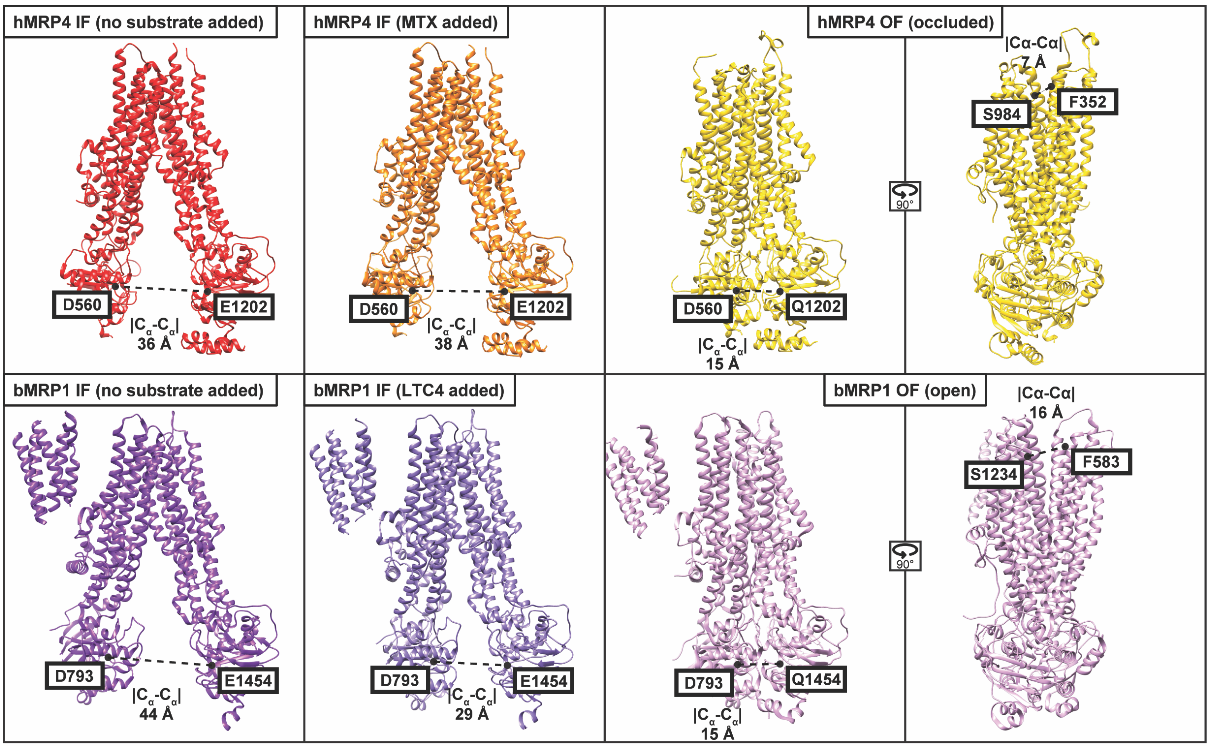

**Fig. S11: Comparison of resolved conformational states of hMRP4 and bMRP1.**
Ribbon representations of indicated structural models displaying the geometric $C_\alpha$-$C_\alpha$ distance ($|C_\alpha$-$C_\alpha|$) between key residues of the NBDs (D560-E/Q1202 for hMRP4; D793-E/Q1454 for bMRP1) and the extracellular gate (S984-F352 for hMRP4; S1234-F583 for bMRP1). The structural models of bMRP1 have been aligned to the structural models of hMRP4, with the unequivocally outlined nanodisc bilayer (Fig. S7) of the hMRP4 reconstructions used as an absolute frame of reference defining the horizontal (membrane) plane.

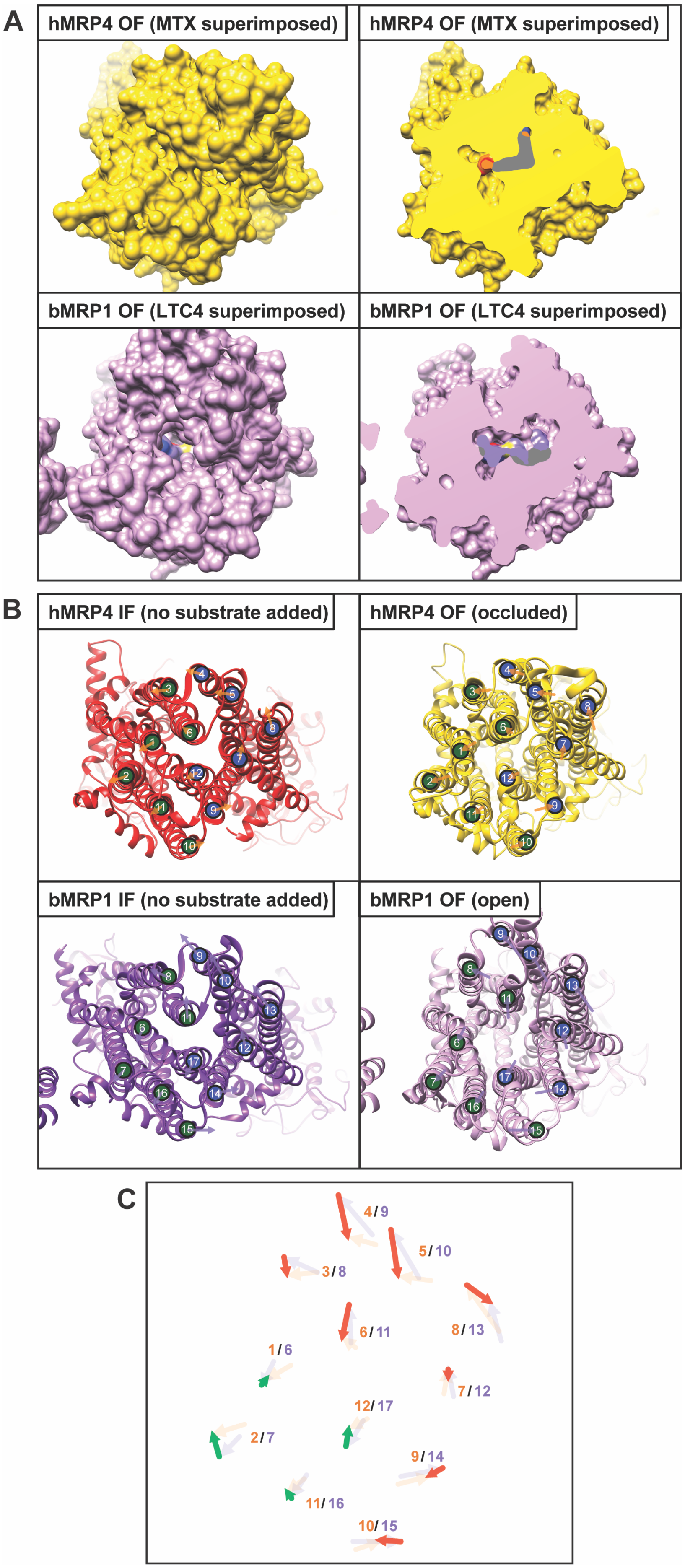
A
hMRP4 OF (MTX superimposed)
hMRP4 OF (MTX superimposed)
bMRP1 OF (LTC4 superimposed)
bMRP1 OF (LTC4 superimposed)
B
hMRP4 IF (no substrate added)
hMRP4 OF (occluded)
bMRP1 IF (no substrate added)
bMRP1 OF (open)
C
4/9
5/10
3/8
6/11
8/13
1/6
7/12
12/17
2/7
9/14
11/16
10/15

**Fig. S12: Conformational rearrangements of hMRP4 and bMRP1.**
(A) Surface representations (viewed from the extracellular side) of indicated structural models of hMRP4 (top row) and bMRP1 (bottom row) onto which surface representations of the MTX (orange) and LTC4 (purple) molecules from the corresponding substrate-bound structural models (aligned to the displayed structural models) have been superimposed. In the slabbed views, gray area indicates steric clash between superimposed substrate and transporter. The structural models of bMRP1 have been aligned to the structural models of hMRP4 as described in Fig. S11.
(B) Ribbon representations (viewed as in (A)) of the indicated structural models of hMRP4 (top row) and bMRP1 (bottom row) onto which TMD helix markers (numbering and color coding according to Fig. 1) have been placed. Markers have been placed at the center of the intersection between the respective transmembrane helix and a common horizontal plane (parallel to the membrane plane) slicing through the middle of the outer bilayer leaflet. Arrows indicate the lateral movement (within the horizontal plane) of respective helices upon IF-OF isomerization of hMRP4 (orange arrows) and bMRP1 (purple arrows).
(C) Enlarged, superimposed view of the movement arrows in (B) with additional arrows (red and green) connecting their endpoints to display the *relative* lateral movement of equivalent transmembrane helices (the identity of the correspondingly moving helix is indicated (orange = hMRP4 transmembrane helix, purple = equivalent bMRP1 transmembrane helix). The additional arrows (red and green) thus display the lateral movement associated with the MRP open-occluded transition, with the movement pattern consisting of two asymmetric elements: An overall rotational motion of 8 transmembrane helices (red arrows) and a predominantly translational motion of the remaining 4 transmembrane helices (green arrows).

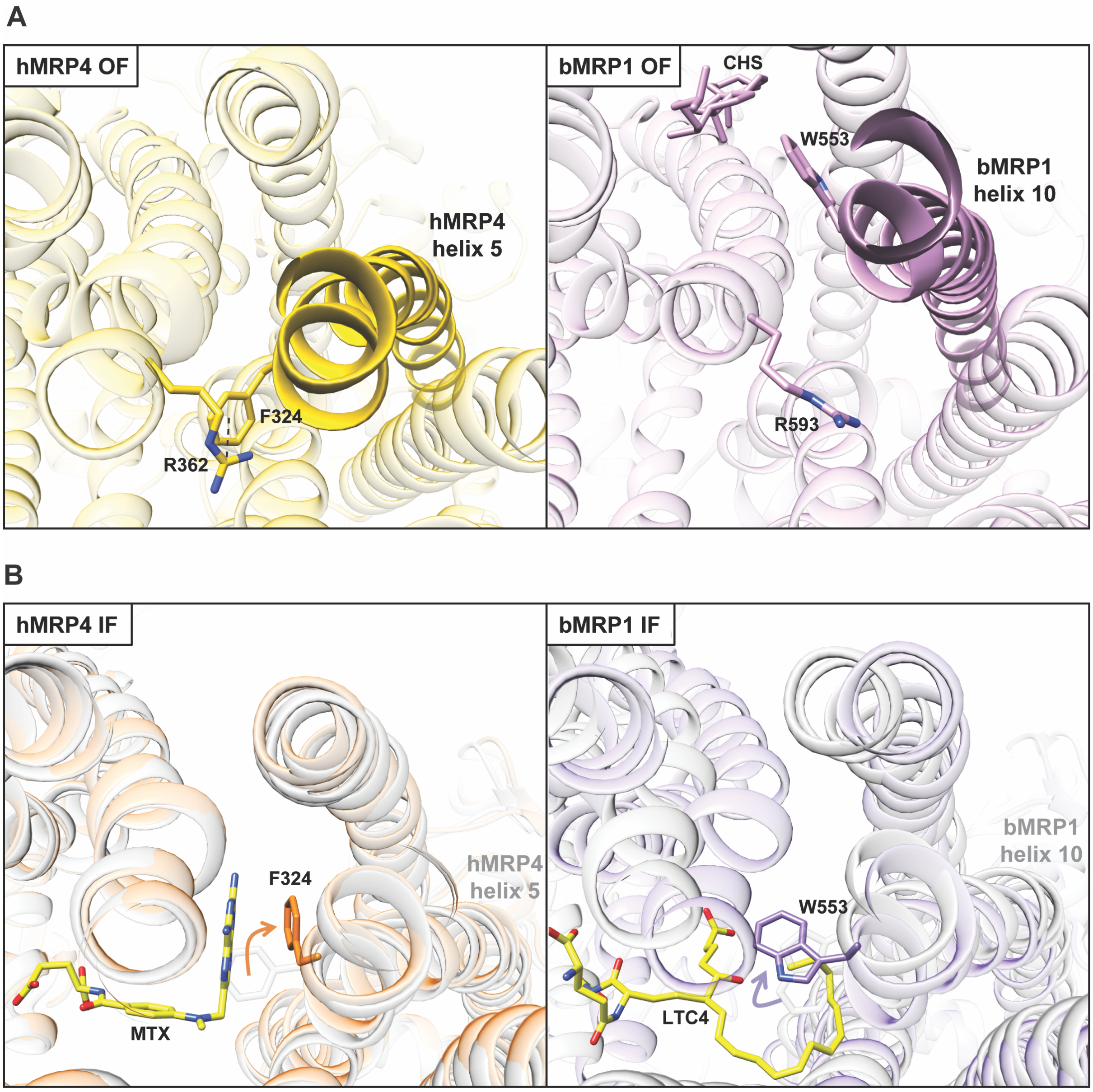

**Fig. S13: Reorientation of equivalent substrate binding residue of hMRP4 and bMRP1.**

(A) Ribbon representations (viewed from the extracellular side) of indicated structural models of hMRP4/bMRP1. Transmembrane helix 5/10 of hMRP4/bMRP1 are highlighted along with equivalent key residues (in the case of bMRP1 OF, also a bound CHS molecule is highlighted). The potential directed cation-$\pi$ interaction between F324 and R362 of hMRP4 is indicated with a punctured line.

(B) Views equivalent to (A) of structural models of IF hMRP4 and bMRP1 (faded orange and purple models represent IF MTX-bound and LTC4-bound structural models of hMRP4 and bMRP1 respectively, while faded gray models represent the respective IF structural models from samples to which no substrate has been added). F324 and W553 of the MTX-bound hMRP4 and LTC4-bound bMRP1 structural models, respectively, are highlighted, and arrows indicate the relative movement of these residues upon substrate-binding. The models of bound MTX and LTC4 molecules are likewise highlighted.

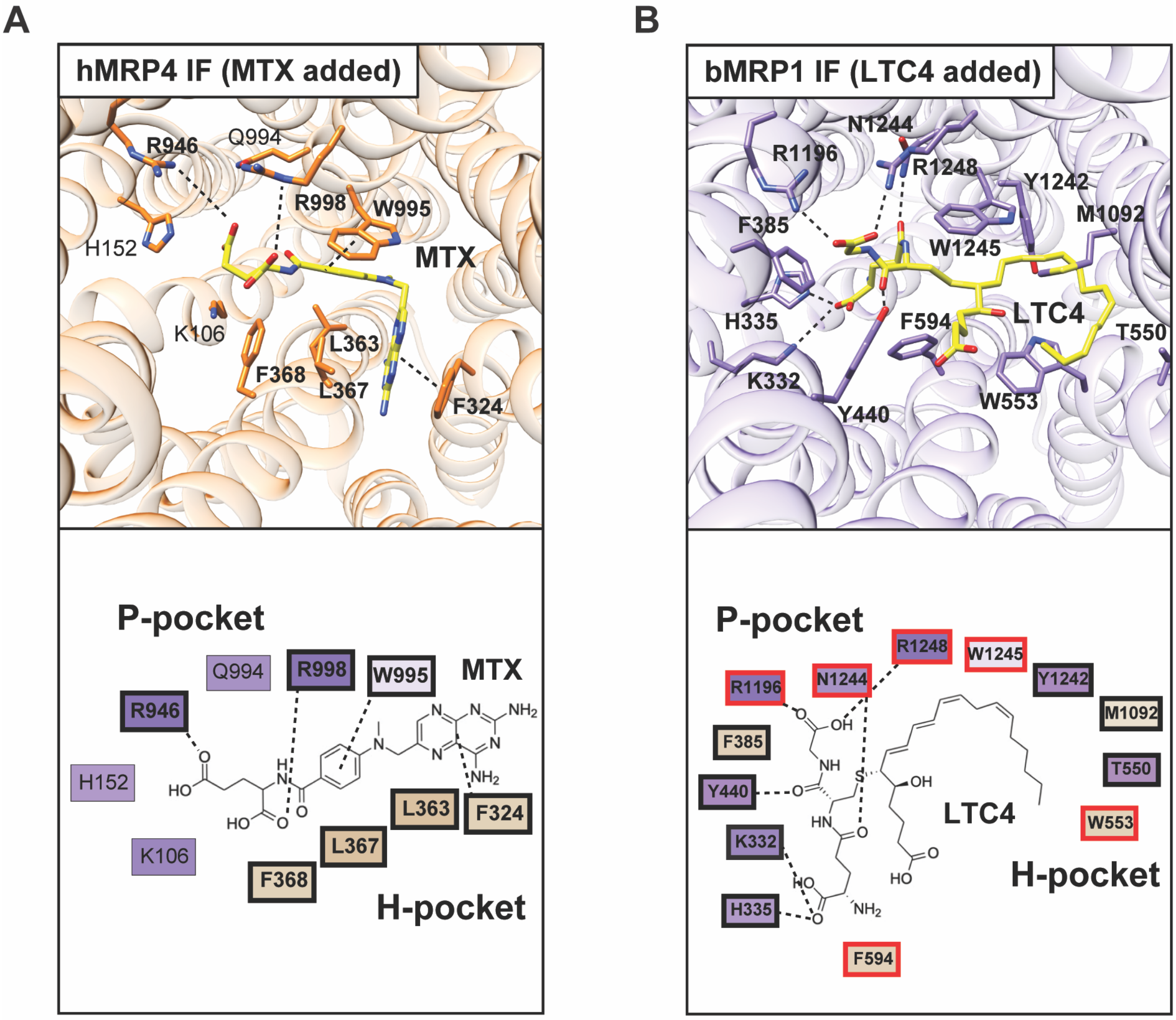

C

| | | | | | | | | | | |
|---|---|---|---|---|---|---|---|---|---|---|
| hMRP7 | 299 G | 341 Q | 510 W | 549 L | - - | 553 N | 1149 Q | 1199 S | 1200 Q | 1203 S |
| hMRP5 | 193 G | 236 L | 406 A | 445 F | - - | 449 V | 1096 R | 1144 Q | 1145 F | 1248 R |
| hMRP8 | 177 S | 220 F | 390 L | 429 L | - - | 433 F | 1044 R | 1092 Q | 1093 A | 1096 R |
| hMRP9 | 137 A | 180 W | 350 A | 389 F | - - | 393 I | 1023 R | 1071 Q | 1072 V | 1075 R |
| **hMRP4** | **106 K** | **152 H** | **324 F** | **363 L** | **367 L** | **368 F** | **946 R** | **994 Q** | **995 W** | **998 R** |
| hMRP6 | 325 R | 367 E | 539 F | 579 K | - - | 583 F | 1169 R | 1217 Q | 1218 W | 1221 R |
| hMRP2 | 336 T | 378 L | 550 F | 591 F | - - | 595 M | 1205 R | 1253 N | 1254 W | 1257 R |
| hMRP1 | 339 M | 381 L | 553 W | 594 F | - - | 598 I | 1197 R | 1245 N | 1246 W | 1249 R |
| hMRP3 | 325 S | 367 L | 539 W | 580 L | - - | 584 M | 1193 R | 1241 N | 1242 W | 1245 R |

**Fig. S14: Comparison of the substrate binding pockets of hMRP4 and bMRP1.** (A) Top: View of the hMRP4 substrate binding pocket (equivalent to the view in Fig. 3) of the structural model of MTX-bound hMRP4. Bottom: 2D representation (based on information presented in Fig. S9 and the ribbon representations above) of MTX coordination with directly interacting residues outlined in bold and directed interactions indicated with punctured lines. Residues are colored according to the Kyte-Doolittle hydrophobicity scale (see Fig. 3).

(B) Top: View of the bMRP1 substrate binding pocket (the structural model of bMRP1 has been aligned to the structural model of hMRP4 as described in Fig. S11) of the structural model of LTC4-bound bMRP1. Bottom: 2D representation of LTC4 coordination[3] with directly interacting residues outlined in bold and directed

interactions indicated with punctured lines. Residues of the bMRP1 substrate binding pocket whose sequence equivalents in the hMRP4 substrate binding pocket interacts with either MTX, TPT or PGE2 are highlighted in red. Residues are colored according to the Kyte-Doolittle hydrophobicity scale (see Fig. 3).
(C) Amino acid sequence alignment of human MRP1-9 showing residues equivalent to 10 key residues of the hMRP4 substrate binding pocket (in bold) colored according to percentage identity. The equivalent *human* MRP1 residues of the *bovine* MRP1 substrate binding pocket whose equivalents in the hMRP4 substrate binding pocket interacts with either MTX, TPT, or PGE2 are highlighted with red boxes.

SUPPLEMENTAL TABLES (TITLES AND LEGENDS)

#### Table S1. Key figures of cryo-EM data collection, processing, refinement, and validation, related to Fig. S2-6.

| | IF MRP4 (no substrate added) | OF MRP4 (TPT added) | IF MRP4 (MTX added) | IF MRP4 (TPT added) | IF MRP4 (PGE2 added) |
|---|---|---|---|---|---|
| **Collection** | | | | | |
| Microscope | Titan Krios G2 | | | | |
| Voltage (kV) | 300 | | | | |
| Magnification (nominal) | 96,000x | | | | |
| Detector | Falcon 3EC | | | | |
| Pixel size (Å) | 0.832 | | | | |
| Defocus range (μm) | -0.5 to -2.5 | | | | |
| Frames/movie | 40 | | | | |
| Dose rate (electrons/pixel/s) | 0.68 | | | | |
| Total exposure dose (electrons/Å$^2$) | 40 | | | | |
| **Processing** | | | | | |
| Movies | 5,391 | 3,378 | 5,032 | 1,449 | 3,071 |
| Number of particles (final) | 725,758 | 90,169 | 382,621 | 56,002 | 107,793 |
| Box size (pixels) | 384 | | | | |
| Map-sharpening B-factor (Å$^2$) | -159.0 | -98.3 | -176.4 | -117.0 | -149.7 |
| Map resolution (Å) (FSC 0.143) | 3.0 | 3.2 | 3.6 | 3.9 | 4.0 |
| **Refinement** | | | | | |
| Model composition | | | | | |
| Non-hydrogen atoms | 9,547 | 9,650 | 9,547 | 9,547 | 9,548 |
| Protein residues | 1,195 | 1,211 | 1,195 | 1,195 | 1,195 |
| Ligands | 6 | 4 | 1 | 1 | 1 |
| RMSZ | | | | | |
| Bond lengths | 0.26 | 0.25 | 0.53 | 0.47 | 0.50 |
| Bond angles | 0.48 | 0.51 | 0.59 | 0.57 | 0.60 |
| CC (mask) | 0.8459 | 0.7853 | 0.8201 | 0.8390 | 0.7822 |
| CC (volume) | 0.8154 | 0.7745 | 0.7940 | 0.8190 | 0.7790 |
| CC (peaks) | 0.5817 | 0.5000 | 0.5957 | 0.5448 | 0.4457 |
| **Validation** | | | | | |
| Molprobity score | 1.24 | 1.32 | 1.34 | 1.41 | 1.66 |
| Clashscore, all atoms | 3.63 | 2.44 | 3.87 | 4.59 | 7.48 |
| Rotamers (%) | | | | | |
| Favored | 96.55 | 97.06 | 97.70 | 98.75 | 95.30 |
| Poor | 0.38 | 0.47 | 0.48 | 0.00 | 0.96 |
| Ramachandran plot (%) | | | | | |
| Favored | 97.56 | 95.76 | 97.05 | 96.97 | 96.38 |
| Allowed | 2.44 | 4.24 | 2.95 | 3.03 | 3.62 |
| Outliers | 0.00 | 0.08 | 0.00 | 0.00 | 0.00 |

#### Table S2. Key figures of cryo-EM data collection and processing, related to Fig. S8 and S9

| | IF MRP4 in native nanodiscs (nothing added) | IF MRP4 in minimal nanodiscs (nothing added) |
|---|---|---|
| **Collection** | | |
| Microscope | Titan Krios G2 | |
| Voltage (kV) | 300 | |
| Magnification (nominal) | 96,000x | |
| Detector | Falcon 3EC | |
| Pixel size (Å) | 0.832 | |
| Defocus range (μm) | -0.5 to -2.5 | |
| Frames/movie | 40 | |
| Dose rate (electrons/pixel/s) | 0.68 | |
| Total exposure dose (electrons/Å$^2$) | 40 | |
| **Processing** | | |
| Movies | 7,335 | 8,513 |
| Number of particles (final) | 242,514 | 175,885 |
| Box size (pixels) | 384 | 384 |
| Map-sharpening B-factor (Å$^2$) | -146.7 | -426.1 |
| Map resolution (Å) (FSC 0.143) | 3.3 | 5.5 |
